# Integrated morphological, multi-omics, and functional profiling reveals microglial plasticity driven by AD risk genes

**DOI:** 10.64898/2026.09.07.749934

**Authors:** Yingjie Zhang, Ai Zhang, Sang Seo, Craig Fredrickson, Min Jung, Xiwei Shan, Lilian Phu, Amber Cramer, Carlos Sanchez-Priego, Vineet Vinay Kulkarni, Christopher M. Rose, Casper C. Hoogenraad, Claire G Jeong

**Affiliations:** Department of Neuroscience, Genentech, Inc., South San Francisco, CA 94080, USA; Department of OMNI Bioinformatics, Genentech, Inc., South San Francisco, CA 94080, USA; Department of Proteomic and Genomic Technologies, Genentech, Inc., South San Francisco, CA 94080, USA

**Keywords:** Microglia, Neurodegenerative diseases, multi-omics, transcriptional states, morphological profiling, gene perturbation, phagocytosis, iPSC-derived models

## Abstract

Microglia, the resident macrophages of the central nervous system, are highly dynamic cells essential for brain homeostasis. While genome-wide association studies (GWAS) strongly implicate microglial dysfunction in Alzheimer’s disease (AD), the mechanistic links coordinating their diverse transcriptional, morphological, and functional states remain poorly understood. Using human induced pluripotent stem cell-derived microglia (iMicroglia) and single-cell RNA sequencing, we identified six distinct transcriptional profiles and mapped them to specific morphological phenotypes via targeted immunofluorescence, establishing a link between microglial morphology and molecular identity. Transcriptomic and morphological profiling further demonstrated profound microglial plasticity, revealing distinct, stimulus-specific responses to AD-relevant pathologies, including Tau PFF, amyloid-beta, and apoptotic neurons. To assess how AD risk variants perturb these states, we performed high-efficiency CRISPR-Cas9 ribonucleoprotein (RNP) knockouts of specific AD GWAS genes. Bulk RNA-seq profiling revealed extensive transcriptional remodeling following genetic perturbation. Crucially, we show that depletion of these AD risk genes disrupts baseline morpho-transcriptomic coupling and fundamentally alters microglial phagocytic capacity. Together, this study reveals how AD GWAS genes may drive microglia into dysfunctional states characterized by altered morphology, distinct multi-omic signatures, and impaired functions.

## Background

Microglia are the resident immune cells of the brain, playing vital roles in development, homeostasis, and neurodegenerative disease progression ^1,2^. During development, microglia eliminate excess synapses via phagocytosis, contributing to neural network formation ^3,4^. In the adult brain, they act as professional phagocytes, clearing debris, dead cells, and pathogens to preserve brain health ^5^. Dysregulated microglial functions are implicated in neurological diseases such as Alzheimer’s disease (AD) and Parkinson’s disease (PD) ^6,7^. However, the limited availability of primary human microglia has hindered mechanistic studies. Recent advances in iPSC-derived-microglia differentiation protocols have achieved varying degrees of success ^8–12^.

Microglia are morphologically and functionally plastic, adopting diverse states across development, aging, and disease ^13,14^. In homeostasis, they exhibit small somas with highly branched processes that support efficient parenchymal surveillance ^15^, whereas pathological activation typically leads to retracted processes and enlarged cell bodies, and ameboid morphology associated with increased phagocytosis and inflammation ^16–18^. Intermediate morphological forms, such as hyper-ramified, and rod-shaped microglia, are linked to specific pathological conditions ^19^ ^20,21^. Morphology often serves as a proxy for microglial state, which is more precisely defined by distinct transcriptional signatures. Single-cell transcriptomics has identified multiple disease-associated states, including disease-associated microglia (DAM) ^22^, microglial neurodegenerative phenotype (MGnD)^23^, proliferative-region-associated microglia (PRAMs) ^24^, and human Alzheimer’s disease microglia (HAMs) ^25^. Yet, directly linking morphological features, transcriptomic identities, and functions remains challenging, underscoring the need for advanced imaging and computational approaches that connect these axes of microglial biology.

Genome-wide AD risk variants preferentially map to microglial enhancers, highlighting the critical roles of genes regulating immune function and phagocytosis ^26–28^. Defining how disease-associated genes shape microglial plasticity–encompassing morphology, dynamic transcriptional states, and core functions–is therefore essential for understanding microglial contributions to pathogenesis. Phagocytosis, a central microglial function, involves coordinated steps of substrate recognition, engulfment, trafficking, and degradation ^29–31^. Impairments in these pathways, often driven by genetic variants, contribute to protein aggregate accumulation, inflammation, and neurodegeneration in AD and PD ^32^. Genome-wide association studies (GWAS) have implicated AD risk genes at multiple nodes of the phagocytic pathways–from substrate recognition (*TREM2*, *complement receptor 1* (*CR1*), *CD33*) to cytoskeletal remodeling during engulfment (*ABI3*, *CASS4*, *PTK2B*), and endolysosomal cargo degradation (*BIN1*, *RIN3*, *ZYX*, *GRN*) ^33^. However, how these genes influence microglial phenotypic and transcriptomic plasticity, as well as phagocytic function, remains poorly understood.

To address these gaps, we optimized an iPSC-derived microglia (iMicroglia) platform, based on the previously reported protocol ^10^, that efficiently generates cells suitable for morphological, transcriptional, and functional analyses. This system recapitulates key features of primary human microglia and enables induction of disease-associated states *in vitro* using stimuli such as amyloid-beta (Aβ) and tau pre-formed fibrils (Tau PFF). To capture the dynamic nature of microglial plasticity, we combined comprehensive morphological profiling with integrated multi-omics. Leveraging this platform alongside Cas9-RNP-based genetic perturbations, we examined how key Alzheimer’s disease (AD) risk genes regulate microglial morphology, transcriptional states, and phagocytosis. Together, this framework provides a new approach for dissecting microglial plasticity, gene function, and their collective contributions to disrupted phagocytic pathways in neurodegenerative diseases.

## Results

### iPSC-derived microglia recapitulate the functional and molecular characteristics of human microglia

Based on a previously published protocol ^10^, we developed a streamlined,reproducible protocol for large-scale generation of mature iPSC-derived microglia (iMicroglia) across three independent iPSC lines (iPS11, KOLF2.1J, and iPS26) (Supplementary Fig. 1A-B). Hematopoietic progenitors were generated by day 10 and matured with interleukin-34 (IL-34), macrophage colony-stimulating factor (M-CSF), and transforming growth factor β (TGF-β)^34,35^, yielding cells expressing homeostatic microglia markers P2RY12 and TMEM119 (Refer to Methods) that were not observed in human monocyte-derived macrophages (hMDM)(Fig. 1A). iMicroglia phagocytosed pHrodo-conjugated synthetic fibrillar amyloid-beta (Aβ-pHrodo) with uptake enhanced by LPS or tau preformed fibrils (Tau PFF) stimulation (Supplementary Fig. 1C-D), and mounted robust cytokine/chemokine responses to LPS and Tau PFFs but not to low-dose Aβ oligomers, consistent with known dose-dependent effects of oligomeric Aβ ^36^ (Supplementary Fig. 1E).

**Figure 1.**
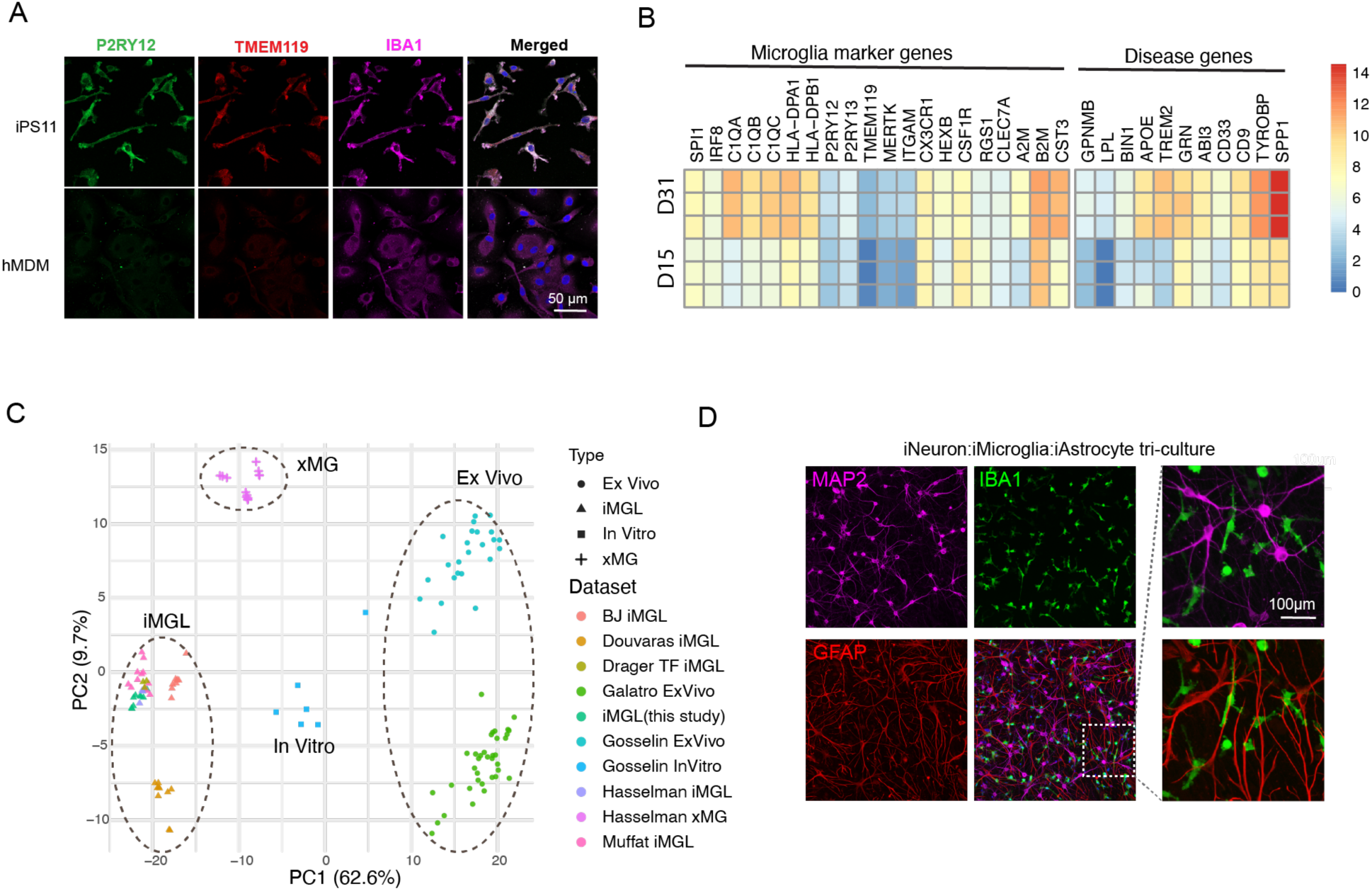
iPSC-derived microglia display functional and transcriptional profiles comparable to human microglia. **A.** Representative immunofluorescence images show robust expression of the homeostatic microglial markers P2RY12 (green), TMEM119 (red), and IBA1 (magenta) in mature iMicroglia (day 31), but not in hMDMs. DAPI (blue) labels nuclei. **B.** Comparison of gene expression levels from the bulk RNA-seq data for canonical microglia markers and disease risk genes between macrophage precursors (D15) and mature iMicroglia (D31). **C.** PCA clustering analysis using the top 2 PCs for our iMicroglia and previously published datasets. PSC-derived microglia like cells (iMGL), xenografted microglia (xMG), *In Vitro* represents primary human microglia cultured *in vitro* in Gosselin et al. **D.** Representative immunofluorescence images of tri-cultures show iPSC-derived microglia labeled with IBA1, iNeurons labeled with MAP2, and iAstrocytes labeled with GFAP. The right panel displays magnified views of the regions highlighted by the boxed areas.

Bulk RNA-seq confirmed the expected transcriptomic profiles of iMicroglia expressing *P2RY12*, *TMEM119*, complement genes such as *C1QA*, *C1QB*, *C1QC*, and AD-associated genes *GPNMB* and *TREM2* (Fig.1C). Proteomics analysis confirmed these markers at the protein level, with a strong correlation (r = 0.664) between RNA and protein changes (Supplementary Fig. 1F). Gene Ontology (GO) analysis of Day 31 iMicroglia showed significant enrichment of immune-related terms (top 5 terms, FDR < 0.05) (Supplementary Fig. 1H). Our iMicroglia transcriptomes clustered closely with other published iPSC-derived microglia datasets and more closely resembled xenografted and *in vitro* human microglia than *ex vivo* samples ^9,37,38^ ^39^ ^40,41^ (Fig. 1C). TGF-beta omission shifted cells toward a peripheral macrophage-like state, confirming its critical role in microglial identity ^42,43^ (Supplementary Fig.1I). In tri-culture with iPSC-derived Ngn2 neurons and iAstrocyte progenitors, iMicroglia adopted a more ramified morphology and distributed evenly among neurons and astrocytes (Fig. 1D), supporting their use for modeling microglia-neuron-astrocyte interactions in disease-relevant contexts. Together, this characterization ensures the quality, reproducibility, and physiologically relevant context of iMicroglia, supporting their utility to interrogate neurodegenerative diseases.

### Single-cell profiling reveals six diverse transcriptional states in iMicroglia

The inherent heterogeneity and dynamic cellular states of human microglia ^44^ prompted us to use single-cell RNA sequencing (scRNA-seq) to characterize this diversity and determine whether distinct microglial states are captured in iMicroglia. We analyzed the single transcriptomes pooled from iMicroglia differentiated from three iPSC lines (iPS11, KOLF2.1J and iPS26), and visualized the data using Uniform Manifold Approximation and Projection (UMAP) (Fig. 2A, supplementary Fig. 2A). Consistent with previous reports ^45,46^, iMicroglia did not separate into clearly distinct clusters representing different sub-types. Instead, they formed a gradient-like continuum of states within two global clusters. Within this continuum, we defined six clusters based on well-established microglial marker genes: antigen-presenting cluster with elevated *HLA class II* expression; a neurodegeneration associated DAM cluster marked by *APOE*, *GPNMB*, and *SPP1;* a lipid-associated cluster expressing elevated *ABCA1*; an inflammatory cluster enriched for *MPO, PRTN3, S100A8*; a cycling cluster expressing cell cycle-related genes; and a homeostatic cluster representing the baseline microglial state from which the other clusters differ by upregulation of specific genes. All clusters expressed comparable levels of *AIF1 (IBA1)*, *CX3CR1* and *P2RY6,* confirming their microglial identity (Fig. 2B). Cluster distributions were largely consistent across the three iPSC lines, although KOLF2.1J-derived iMicroglia exhibited a proportionally larger homeostatic cluster and a slightly more prominent inflammatory cluster (Supplementary Fig. 2B). Together, these results demonstrate that iMicroglia recapitulate the complex, continuous transcriptional landscape characteristic of human microglia.

**Figure 2.**
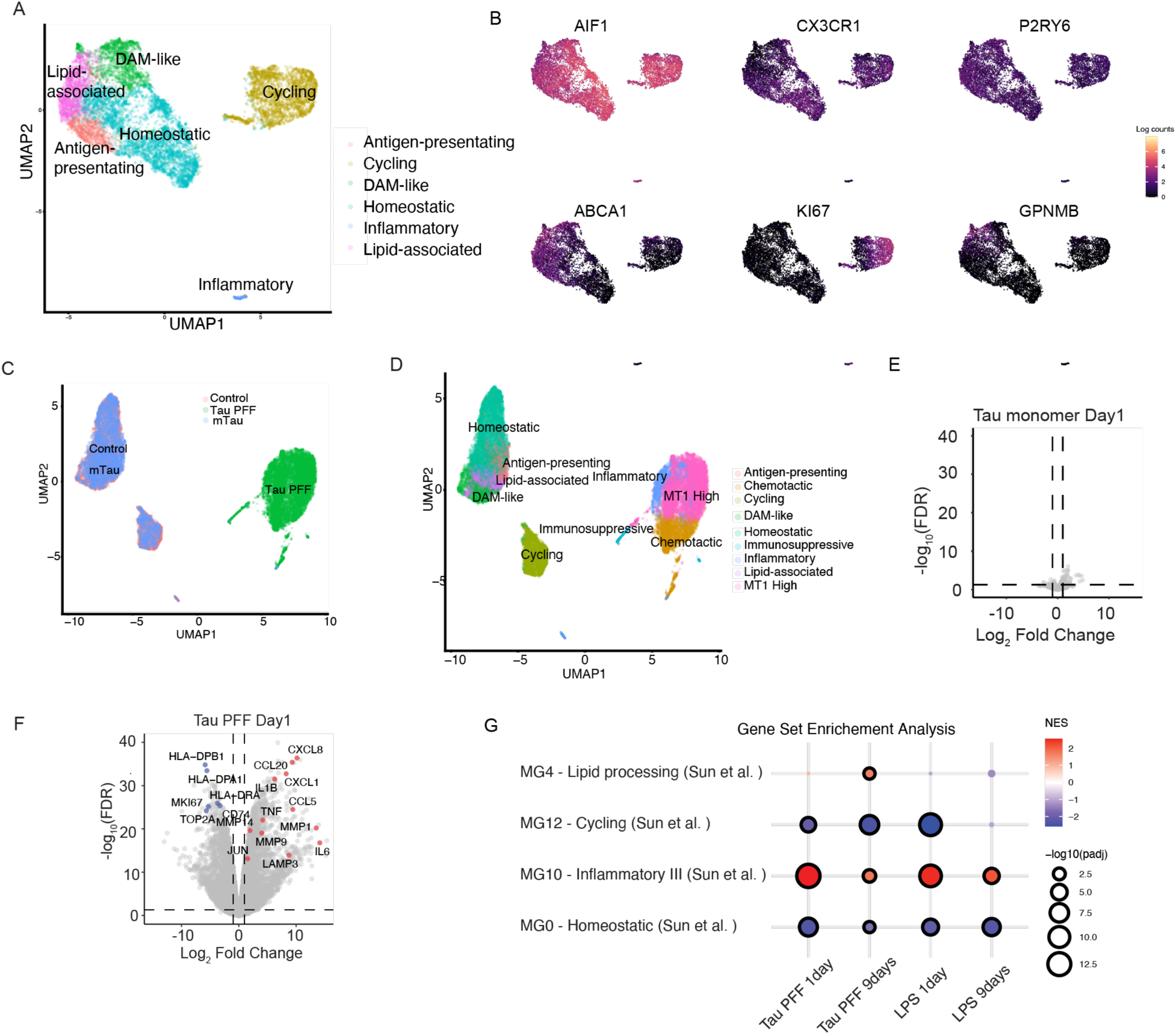
Single-cell transcriptomic analysis reveals six distinct iMicroglia states that undergo dynamic transitions toward inflammatory or oxidative stress– adaptive states in response to Tau PFF stimulation. **A.** UMAP analysis of mature iMicroglia reveals six clusters: antigen-presenting, cycling, DAM-like, homeostatic, inflammatory and lipid-associated clusters. **B.** UMAP shows the selected marker genes expressed in the different states identified in iMicroglia. *ABCA1*, *KI67* and *GPNMB* are highly enriched in the lipid-associated, cycling and DAM-like clusters respectively. *CX3CR1* is more enriched in homeostatic,antigen-presenting and cycling clusters. All clusters express similar levels of *AIF1* and *P2RY6*. **C.** UMAP projection of single-cell data demonstrates that Tau PFF-treated cells form a distinct cluster, clearly separate from untreated control and Tau monomer-treated cells. **D.** Nine distinct microglia states were defined through clustering analysis in Fig. 2C. Tau PFF-treated cells exhibited unique chemotactic, immunosuppressive and *MT1*-High states, along with a minor population of cycling cells that overlapped with control and Tau monomer populations. **E.** Volcano plot of differential expression analysis for Tau monomer treatment, showing minimal transcriptional change relative to control. **F.** Volcano plot of differential expression analysis results from Tau PFF exposure. Tau PFF treatment induced substantial gene expression changes, including upregulation of inflammatory response genes and downregulation of HLA class II complex and cell cycle genes. Selected genes from relevant pathways are highlighted. **G.** GSEA of bulk RNA-seq shows significant upregulation of inflammatory III pathway genes at day 1 following Tau PFF or LPS treatment, corresponding to the MG10 cluster in the *in vivo* dataset reported by Sun et al. Notably, only prolonged Tau PFF-treated iMicroglia (day 9) exhibit significant enrichment of the lipid processing (DAM-like) signature associated with the MG4 cluster from the same dataset, distinct from prolonged LPS treatment (presence of dotted outlines: adjusted p value < 0.05).

### Acute Tau PFF exposure shifts iMicroglia toward more inflammatory and oxidative stress–adaptive states

To investigate how iMicroglia respond to brain-relevant stimuli, we performed scRNA-seq on naive and Tau PFF- or monomer-treated iMicroglia derived from iPS11 and KOLF2.1J, yielding a dataset of 17,374 single cells across three biological replicates (Fig. 2C). Dimensionality reduction and clustering analysis showed that Tau monomer-treated cells clustered with untreated controls (Fig. 2C), suggesting minimal effects on iMicroglia states, whereas Tau PFF-treated cells formed a distinct cluster, indicating that Tau PFF exposure induces unique transcriptional states (Fig. 2C). UMAP analysis identified nine microglial states overall (Fig. 2D, supplementary Fig. 2C): Tau monomer-treated cells mapped onto the six states observed in untreated controls (Fig. 2D, supplementary Fig. 2C), whereas Tau PFF-treated cells showed minimal overlap with untreated control states (Fig. 2D, supplementary Fig. 2C). Tau PFF exposure was predominantly enriched in three state clusters: *Metallothionein-1*(*MT1*)-high, Inflammatory, and chemotactic clusters, demonstrating that, unlike Tau monomers, Tau PFFs elicit inflammatory activation and oxidative-stress adaptive transcriptional shifts in iMicroglia (Supplementary Fig. 2C). To characterize these states further, we examined cluster-specific marker genes. In the immunosuppressive cluster of Tau PFF-treated cells, *EBI3*, *CCL22* and *NFKBIA* were upregulated, while the chemotactic cluster expressed high levels of *CCL2*, *CCL3*, *CXCL8*, *S100A8* and *S100A9*. The *MT1*-high cluster was strongly enriched for metallothionein family genes, including *MT1G, MT2A, MT1H, MT1M and MT1X, MT1F* and *MT1E*. Dysregulation of *MT1* has been associated with neurodegeneration ^47,48^. In AD, upregulated MT proteins may function as a protective mechanism against oxidative stress and cytokine exposure, thereby reducing neuroinflammation ^49–51^. Thus, the emergence of a predominant *MT1*-high cluster in Tau PFF-treated iMicroglia may reflect a metallothionein-mediated immunomodulatory and adaptive neuroprotective response. These findings highlight the complex, state-specific inflammatory responses elicited by Tau PFF exposure.

### Prolonged Tau PFF exposure models disease progression

To assess the temporal dynamics of these responses, we conducted bulk RNA-seq on iMicroglia treated with soluble Tau monomers or insoluble Tau PFFs for 1, 5, and 9 days. While Tau monomers caused minimal transcriptional changes (Fig. 2E), differential expression analysis of Tau PFF-treated with control cells identified 6,619 genes with significant changes (3,232 upregulated and 3,387 downregulated;FDR < 0.05, log2 Fold Change >1 or < -1) (Fig. 2F). Notably, inflammatory markers such as *TNFa*, *IL-6*, *IL-1*, *CXCL8* and *CCL-2* were upregulated in Tau PFF-treated iMicroglia (Fig. 2F). KEGG pathway analysis indicated that Tau PFF exposure upregulated pathways associated with inflammatory responses, including cytokine-cytokine receptor interaction, *TNF*, *IL-17* and *NF-kappa B* signalling pathways (Supplementary Fig. 2D), while downregulated pathways mainly associated with cell cycle and DNA replication (Supplementary Fig. 2E).

We contextualized these transitions using *in vivo* microglial gene signatures described by Sun et al. ^52^. Gene set enrichment analysis (GSEA) showed acute inflammatory signatures corresponding to MG10 state (Sun et. al.), which were significantly elevated at all time points, peaking at day 1 and gradually declining by days 5 and 9 (Supplementary Fig. 2F). By day 9, Tau PFF-treated cells additionally showed enrichment of the *in vivo* MG4 signature, capturing canonical DAM and MGnD state markers, such as *APOE*, *ABCA1*, and *TREM2*. To verify the specificity of this temporal shift, we compared Tau PFF treatment to LPS exposure. Both treatments induced comparable inflammatory signatures at day 1; however, by day 9, the DAM-like enrichment was more upregulated in the Tau PFF-treated samples (Fig. 2G). Sun et al. identified three distinct inflammatory clusters (MG10, MG2, and MG8) and proposed a progression model in which the inflammatory response is most potent in early stages (MG10) before dampening or transforming over the disease course (MG10 > MG2 > MG8), with MG10 representing an early, acute inflammatory state and MG8 a chronic, end-stage phenotype. Our data showed a similar trend: acute Tau PFF treatment drives cells toward the MG10-like signature, whereas prolonged exposure shifts the phenotype away from acute cytokine expression toward a more upregulated canonical DAM-like profile. Although this warrants further investigation using diverse CNS-relevant stimuli (i.e. Aβ plaques, distinct Tau isoforms, and varied fibril structures), this highlights the potential utility of the iMicroglia model for studying the dynamic evolution of the inflammatory cascade and slow disease progression in AD.

### Distinct pathological stimuli elicit shared and stimulus-specific transcriptional signatures in iMicroglia

Given the transcriptional alterations observed with Tau PFFs, we further characterized iMicroglia plasticity by exposing them to other brain-relevant pathological stimuli, including Aβ fibrils and apoptotic neurons (ANs) via bulk RNA-seq. All treatments induced transcriptomic changes, with 445 genes commonly upregulated across conditions (Supplementary Fig.3A). GO analysis of these 445 genes revealed enrichment in vesicle organization, transport, and macroautophagy pathways (Supplementary Fig. 3B), suggesting a shared biological response. Despite this overlap, each stimulus also produced distinct transcriptional signatures. Tau PFFs induced a broad array of inflammatory pathways, including chemotaxis, IL-1, IFN-γ, NF-κB, TNFα, and suppressed antigen-presentation and cell-cycling pathways (Supplementary Fig. 3C). In contrast, Aβ fibrils selectively enhanced lipid- and metabolism-related pathways while downregulating small GTPase and adhesion-related genes, while ANs elicited no statistically significant pathway changes. Both Aβ fibrils and ANs elicited a more modest shift toward an *APOE*/*GPNMB*-high (DAM-like) state compared to the pronounced inflammatory phenotype driven by acute Tau PFF exposure (Supplementary Fig. 3D), noted by strong upregulation of inflammatory genes (*IL6, IL1B, CXCL5*) and downregulation of antigen-presentation and cell cycle markers (Supplementary Fig. 3D). These findings demonstrate that iMicroglia are highly plastic and capable of mounting distinct, stimulus-dependent state transitions that mirror the adaptive nature of microglia across diverse pathological environments in *vivo* ^52,53^.

**Figure 3.**
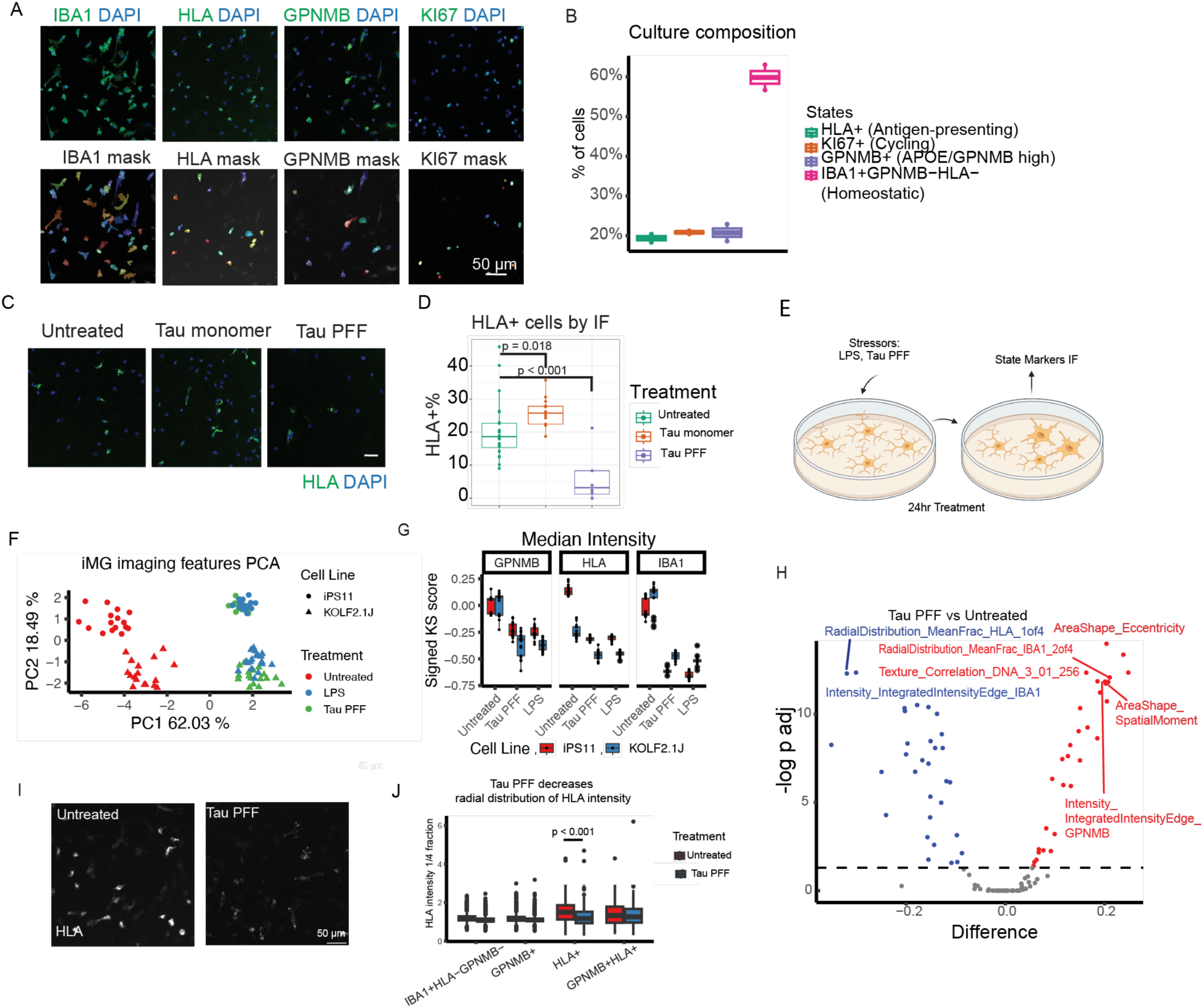
Distinct morphological and transcriptional shifts in iMicroglia reflect stimulus-specific responsiveness to CNS disease-relevant cues. **A.** Top cell state markers which showed a binary expression pattern are used to indicate cell state with IF staining (Antigen-presenting: HLA-DRA, Cycling: KI67, APOE/GPNMB high (DAM-like): GPNMB, Homeostatic: IBA1+HLA-GPNMB-. Cellpose segmentation models were trained to recognize HLA+, KI67+, GPNMB+ and IBA1+ cells. **B.** Quantification of the percentage of cells in each state in iMicroglia cultures, as determined by IF imaging analysis in (A), reveals a state composition consistent with that identified by scRNA-seq analysis in Fig. 2A. **C.** IF imaging analysis confirmed Tau PFF treatment-induced decrease in HLA markers. **D.** Barplot showed the percentage of the HLA+ populations was reduced in Tau PFF-treated cells, consistent with the observations in scRNA-seq data. **E.** Experimental schematic illustrating iMicroglia treatment with LPS or Tau PFFs for 24 hours, followed by IF staining with antibodies against IBA1, GPNMB, and HLA. **F.** PCA of iMicroglia morphological features at the well level. Shapes indicate cell lines and colors indicate treatments. Tau PFF and LPS treatments appear to be similar in the morphology feature space. The two cell lines iPS11 and KOLF2.1J are separated on PC2. **G.** GPNMB, HLA and IBA1 median intensity are significantly decreased by either Tau PFF or LPS treatment. **H.** Volcano plots of Tau PFF treatment vs untreated were shown. Features that were significantly upregulated (red) and downregulated (blue) in treated versus untreated cells are highlighted. The top features were labeled by text. **I.** Representative IF images of HLA staining in untreated and Tau PFF treated cells. HLA localization changes from throughout the cell body to a punctated pattern upon Tau PFF treatment. **J.** Boxplot showed HLA intensity radial distribution in untreated and Tau PFF treated cells, grouped by iMicroglia states. Tau PFF only induces HLA intensity radial distribution changes in HLA+ populations.

### Morphological profiling confirms diverse transcriptomic states and captures stimulus-induced shifts

To confirm the presence of distinct clusters identified by scRNA-seq and to relate transcriptional states to morphological features, we performed immunofluorescence staining for key markers of major microglial state clusters: HLA-DRA for antigen-presenting, KI67 for cycling, GPNMB for DAM-like, and IBA1+HLA-GPNMB- for the baseline homeostatic states (Fig. 3A). Other remaining clusters, such as a lipid-associated cluster expressing elevated ABCA1, could not be confirmed with this approach due to the lack of validated antibodies. We developed a streamlined three-plex IF assay (IBA1, GPNMB and HLA-DRA) to link specific morphological features of iMicroglia to their underlying transcriptional states. CellPose segmentation models trained to recognize HLA+, KI67+, GPNMB+, and IBA1+ cells (described in methods) confirmed the composition of transcriptional states in mature iMicroglia (Fig. 3B). IF staining further demonstrated a reduction in HLA expression following Tau PFF exposure compared with Tau monomers (Fig. 3C-D), consistent with the loss of antigen-presenting cells observed in the scRNA-seq data (Supplementary Fig. 2C). To check if this IF assay can differentiate between different myeloid cell types, we compared iMicroglia and macrophage-like cells, hMDM via bulk RNA-seq and IF staining. hMDM displayed a slight decrease in the microglial marker *IBA1* and *HLA* expression, but a pronounced increase in *GPNMB*, a DAM-like state marker (Supplementary Fig. 4A). IF staining similarly showed decreased proportions of HLA+ or IBA1+ cell populations in hMDM than iMicroglia, yet GPNMB+ cells were more abundant in hMDM (40% vs. 20%) (Fig. 3B, supplementary Fig. 4B), differentiating transcriptional identity between iMicroglia and hMDM.

Microglia dynamically adjust their shape and spatial organization in response to diverse stimuli, and this morphological plasticity plays a critical role in both physiological and pathological processes ^1,54,55^. To relate transcriptional states to morphological plasticity in iMicroglia, we first profiled morphological features in response to brain-relevant stimuli. iMicroglia were exposed to Tau PFFs or LPS for 24 hours and stained with antibodies against IBA1, GPNMB, or HLA to capture state-associated morphological signatures (Fig. 3E). Segmentation models for IBA1+, GPNMB+, and HLA+ cells were trained using Cellpose2, and applied in CellProfiler to identify labeled cells. We extracted ∼400 single-cell features–including size, shape, intensity distribution and texture from IBA1, GPNMB, HLA and DAPI channels (Supplementary Fig. 4D, E). HLA and GPNMB signals showed minimal overlap (Supplementary Fig. 4F), consistent with our scRNA-seq evidence that they mark distinct transcriptional states. Individual iMicroglia were visualized using PHATE algorithm ^56^, and thumbnails of 10 randomly selected cells from each Cellpose-determined state were placed at their corresponding coordinates (Supplementary Fig. 4G). Cells with high GPNMB or HLA intensity coincided with Cellpose-determined GPNMB+ or HLA+ states, respectively, confirming the segmentation accuracy (Supplementary Fig. 4H).

To assess the overall effect of acute LPS and Tau PFF treatment on morphological features, single-cell level features were aggregated to well-level features using signed KS normalization and analyzed by PCA. Untreated iMicroglia separated from LPS- and Tau PFF-treated cells along PC1 (Fig. 3F), while PC2 captured donor-iPSC line differences (Fig. 3F). To relate these morphological signatures to transcriptional states, we assessed expression of microglial state markers (Fig. 3G). All markers were downregulated in LPS- or Tau PFF-treated cells, with substantial decreases in both IBA1 and HLA, suggesting reduced homeostatic and antigen-presenting states (Fig. 3G). Extending this analysis to fibrillar Aβ and apoptotic neurons (ANs) exposure revealed that PC1 captured a general treatment effect across all three lines, although the shift was much milder than observed with LPS or Tau PFFs, while PC2 distinguished Aβ- and ANs-induced responses (Supplementary Fig. 4I). Among state markers, GPNMB showed a modest increase and IBA1 a decrease, indicating a subtle shift from homeostatic to a more DAM-like phenotype with both treatments (Supplementary Fig. 4J).

To identify the features most affected by treatment, we performed one-way ANOVA with post hoc Tukey testing, focusing on Tau PFF given its similarity to the LPS response (Fig. 3H). Tau PFF-treated iMicroglia generally exhibited a more rod-like shape (increased eccentricity), reduced IBA1 intensity, and enhanced GPNMB edge intensity. A key feature reduced by Tau PFF exposure was HLA intensity in the innermost 25% of the cell (RadialDistribution_MeanFrac_HLA_1of4, Fig. 3H), suggesting a change in the subcellular localization of HLA protein. Indeed, Tau PFF exposure not only lowered overall HLA intensity but also shifted its spatial pattern from diffuse to punctate (Fig. 3I– J), indicating an altered state of the HLA+ population. Collectively, LPS and Tau PFFs induced profound morphological remodeling, whereas fibrillar Aβ and ANs elicited comparatively subtle changes. These data show that we can capture diverse morphological features of iMicroglia and relate them to their transcriptional states, providing insight into how microglia integrate external stimuli into coordinated structural and molecular responses.

### Depletion of *TREM2*-associated AD GWAS genes alters microglial morphology

While multi-omics and morphological profiling of iMicroglial have mapped the broad spectrum of microglial states and their interrelationships, the genetic drivers mediating these transitions and their functional consequences remain largely untested. To address this gap, we selected a panel of Alzheimer’s Disease (AD) GWAS genes known to influence microglial morphology, transcriptional state, or phagocytosis, implicated in *TREM2* signaling in particular, given *TREM2*’s established role in controlling microglial state transitions ^57,58^. We performed CRISPR-Cas9 ribonucleoprotein (RNP) knockouts of the selected targets with an optimized Cas9/RNP editing protocol (see Methods), and evaluated the resulting morphological changes. All target-gene RNPs achieved >50% mRNA knockdown (Supplementary Fig. 5A), and IF imaging confirmed the absence of RAB10, SHIP1, and TREM2 protein expression in the corresponding knockout cells (Supplementary Fig. 5B–D).

We first examined bright-field morphology of the knockout iMicroglia. While *RAB10* and *TREM2* knockout cells resembled controls, depletion of *PTPN6* or *INPP5D* produced striking morphological changes, yielding enlarged, rounded, donut-shaped cells (Fig. 4A). Co-depletion of *INPP5D* and *PTPN6* exacerbated these effects, producing a more severe phenotype than either single knockout. Quantification of brightfield features, such as cell area, radius, eccentricity, confirmed this: cumulative distribution analyses revealed a pronounced rightward shift in cell area and radius for *PTPN6*- and/or *INPP5D*-deficient cells, indicating globally increased cell size, alongside a leftward shift in eccentricity, confirming a rounder morphology (Fig. 4B). PCA of all bright-field features further showed that *PTPN6* and *INPP5D* knockouts clustered separately from controls along PC1, with the double knockout even more distinct, whereas *RAB10* and *AURKB* knockouts aligned more closely with controls along PC2, consistent with subtler morphological effects (Fig. 4C).

**Figure 4.**
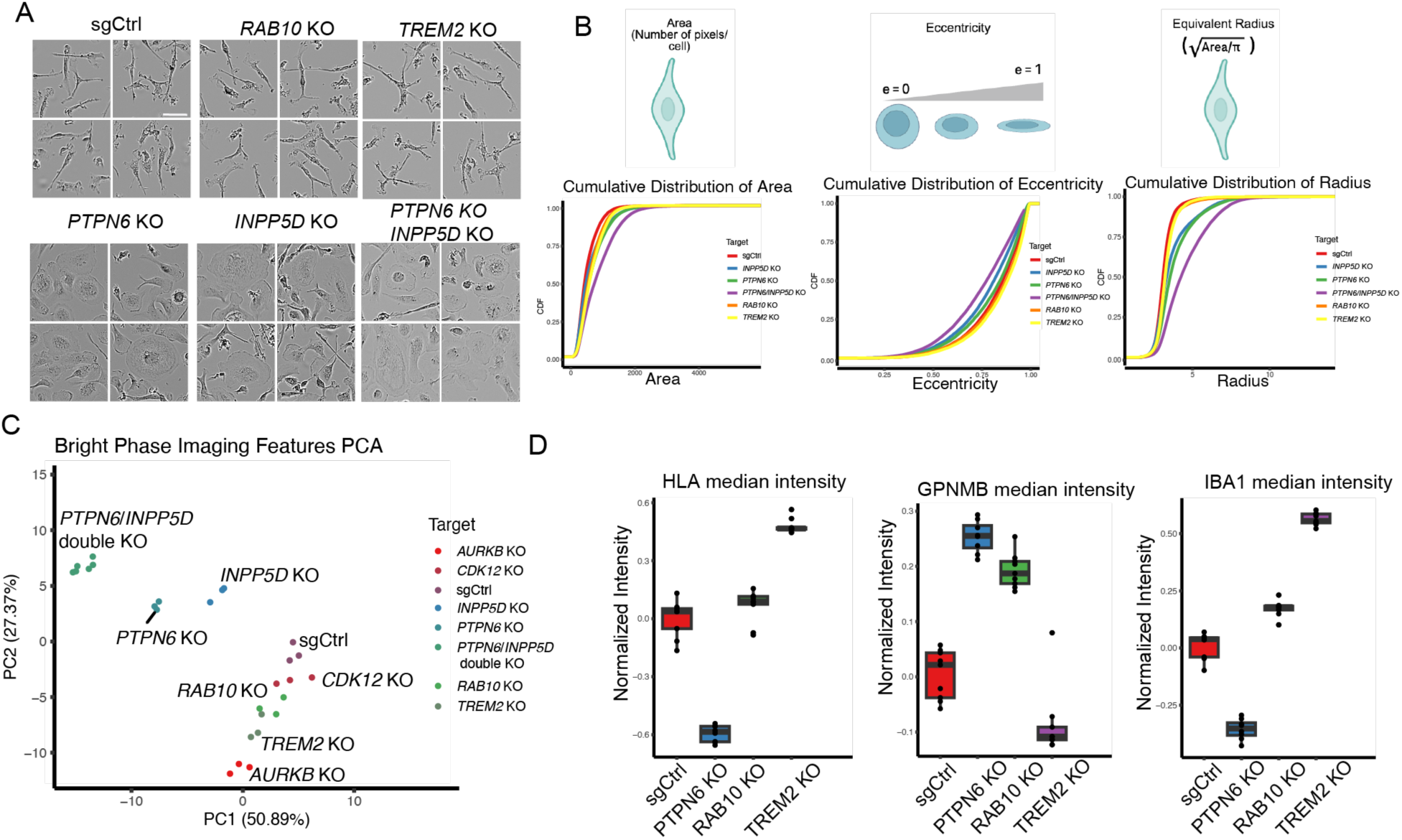
Morphological profiling links microglial state markers to distinct cellular phenotypes and stimulus-induced remodeling. **A.** Bright field images of iMicroglia expressing control, *RAB10*, *TREM2*, *PTPN6*, *INPP5D* sgRNAs. Scale bar, 50 µm. **B.** Representative morphological profiling features such as Area, Eccentricity and Radius definitions were illustrated. Cumulative distribution analysis of these parameters in iMicroglia reveal shifts in cellular morphology in *PTPN6* and *INPP5D* knockout cells compared with controls. **C.** The PCA plot of brightfield imaging features shows a correlation between PC1 and donut-shape morphology (*PTPN6* KO, *INPP5D* KO and *PTPN6*/*INPP5D* double KO), and between PC2 and homeostatic morphology (*AURKB* KO and *RAB10* KO). **D.** Box plots of median intensity for HLA, GPNMB, and IBA1 in iMicroglia expressing control, *RAB10*, *PTPN6* and *TREM2* sgRNAs (n=9 images). The observed changes in HLA, GPNMB, and IBA1 levels point to shifts in microglial states.

We next performed morphological profiling based on IF staining of state markers (IBA1, HLA, GPNMB). This analysis revealed clear state transitions in *PTPN6*- and *INPP5D*-deficient cells (Fig. 4D). *PTPN6* knockout reduced homeostatic features, while *TREM2* depletion decreased GPNMB (DAM-like) and increased IBA1 expression, consistent with reports that TREM2 depletion suppresses GPNMB and APOE expression in microglia ^32,59^.

### CRISPR-Cas9 Knockout of AD GWAS genes drives extensive transcriptomic remodeling

To further probe how these AD GWAS genes regulate microglial states, we examined their transcriptomic effects by bulk RNA-seq. PCA of RNA-seq data revealed substantial gene expression changes following deletion of each gene (Fig. 5A). *RAB10* deficiency produced the most extensive transcriptional changes, with 2,999 genes upregulated and 2,516 downregulated, including major ITAM and ITIM signaling molecules such as *TREM2*, *CD33*, and *PTPN6* (Fig. 5B); GSEA analysis further showed increased lysosomal activity in *RAB10*-deficient iMicroglia (Supplementary Fig. 6A). Both *INPP5D* and *PTPN6* knockouts induced distinct transcriptomic shifts (Fig. 5C–D), with enhanced stress-response pathways in *INPP5D* knockouts (Supplementary Fig. 6B) and enriched extracellular matrix–related pathways in *PTPN6* knockouts (Supplementary Fig. 6C).

**Figure 5.**
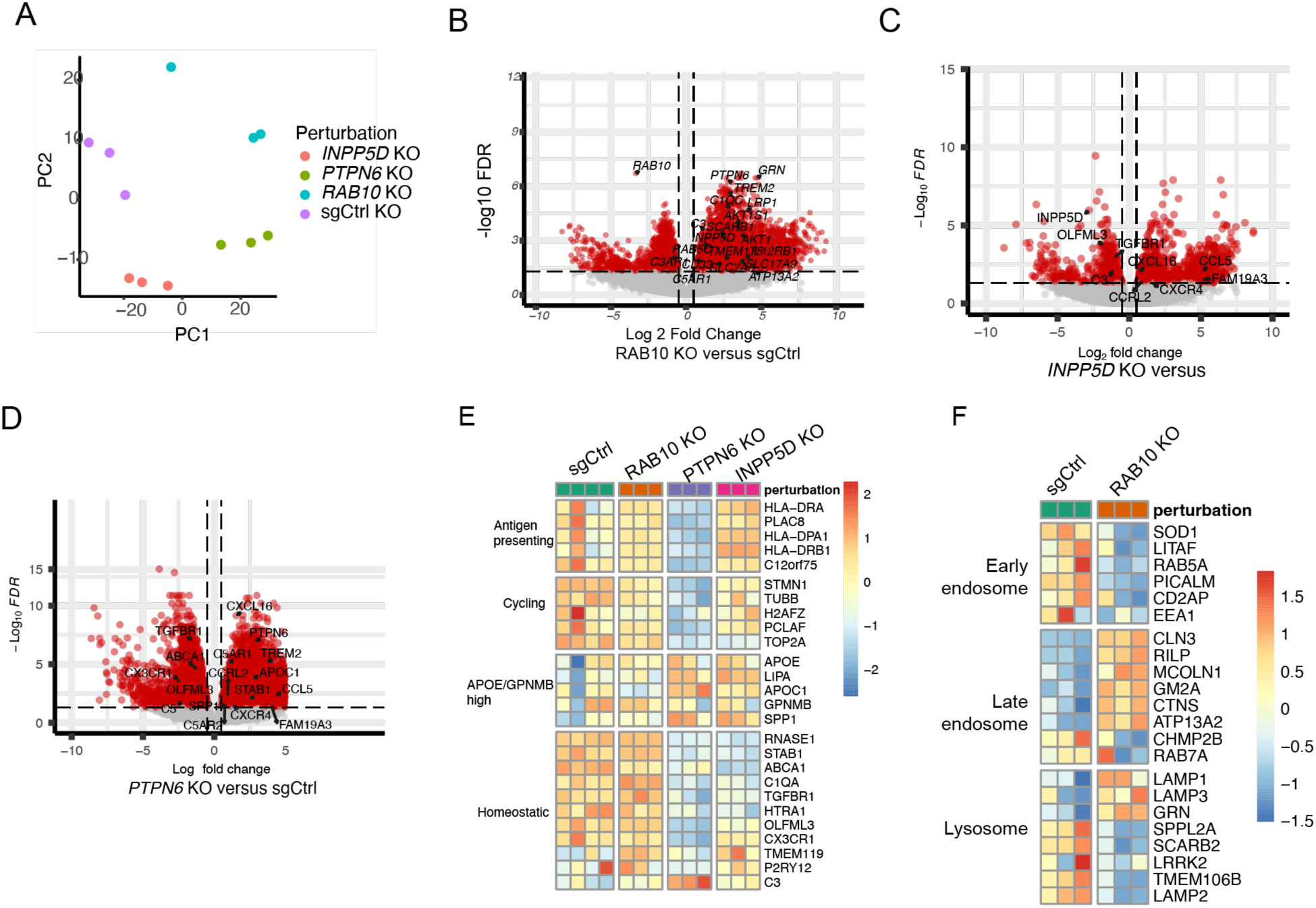
CRISPR-Cas9 KO of AD GWAS genes drives transcriptional state changes. **A.** PCA of RNA-seq datasets from iMicroglia expressing control, *INPP5D*, *PTPN6*, or *RAB10* sgRNAs. The PCA plot showed that *RAB10*, *PTPN6,* and *INPP5D* knockout cells clustered separately from control cells, with PC1 accounting for 22.31% and PC2 for 12% of the variation, suggesting substantial transcriptome alterations. **B.** Volcano plot of differential gene expression from RNA-seq data in *RAB10* knockout versus control iMicroglia (n=3). Red dots indicate statistically significant upregulated (2,999) and downregulated (2,516) genes (log2 fold change > 0.5, FDR < 0.5). **C.** Volcano plot showing the differential gene expression profile of *INPP5D* KO versus control (sgCtrl) iMicroglia as determined by RNA-seq (n = 3). Genes with log_2_ Fold change >0.5, FDR < 0.5 are shown. Statistically significant 989 upregulated and 524 downregulated genes are indicated in red dots. **D.** Volcano plot showing the differential gene expression profile of *PTPN6* KO versus control (sgCtrl) iMicroglia as determined by RNA-seq (n = 3). Genes with log_2_ Fold change >0.5, FDR < 0.5 are shown. Statistically significant 5,295 upregulated and 3,379 downregulated genes are indicated in red dots. **E.** Heatmap of iMicroglia state marker expression across RNA-seq samples (control, *RAB10*, *PTPN6*, and *INPP5D* sgRNAs). *PTPN6-* and *INPP5D*-knockout iMicroglia showed downregulation of homeostatic and upregulation of DAM-like markers compared to control (n=3). **F.** Heatmap of endolysosomal pathway gene expression in iMicroglia (control and *RAB10* knockout sgRNAs)(n=3). *RAB10* knockout iMicroglia displayed dysregulation of numerous endolysosomal genes.

We then examined how these genetic perturbations influenced microglial transcriptional states. RNA-seq analysis showed that *PTPN6* and *INPP5D* knockouts downregulated homeostatic markers while concurrently upregulating DAM-associated genes (Fig. 5E); *PTPN6* depletion additionally reduced antigen-presenting and cell-cycle–related genes. In contrast, *RAB10* knockout had minimal effect on microglial state markers (Fig. 5E). Pathway-level analysis confirmed dysregulation of endosomal and lysosomal genes and activities in *RAB10* knockout cells, consistent with endosomal trafficking inefficiency (Fig. 5F) and pointing to compromised endosomal-lysosomal function.

### Targeted depletion of AD GWAS genes modulates microglial phagocytic capacity in a substrate-specific manner

Given that AD risk gene depletion altered both morphology and transcriptional state, we next investigated the functional significance of this coupling, focusing on phagocytosis as a canonical microglial function critical for brain homeostasis ^60,61^. We performed phagocytosis assays using pHrodo-labeled Aβ fibrils, apoptotic neurons, and zymosan particles, quantifying fluorescent uptake over a 24-hour time course. *AURKB* and *CDK12* knockouts served as controls, since *AURKB* depletion is reported to promote phagocytosis while *CDK12* depletion decreases it ^39,62^; our data were consistent with these prior findings (Fig. 6A–C, Supplementary Fig. 7A–B). Among the AD risk genes tested, *RAB10* knockout enhanced Aβ uptake, whereas *PTPN6* and *INPP5D* knockouts reduced it, as shown by quantification and visualization of intracellular Aβ-pHrodo intensity (Fig. 6A–C). *INPP5D*, a known AD risk gene ^63,64^, significantly decreased Aβ uptake but had minimal effect on apoptotic neuron or zymosan phagocytosis (Fig. 6A– C). *PTPN6*, also implicated in AD ^65^, reduced uptake of both Aβ and apoptotic neurons but not zymosan, suggesting substrate specificity (Fig. 6A–C).

**Figure 6.**
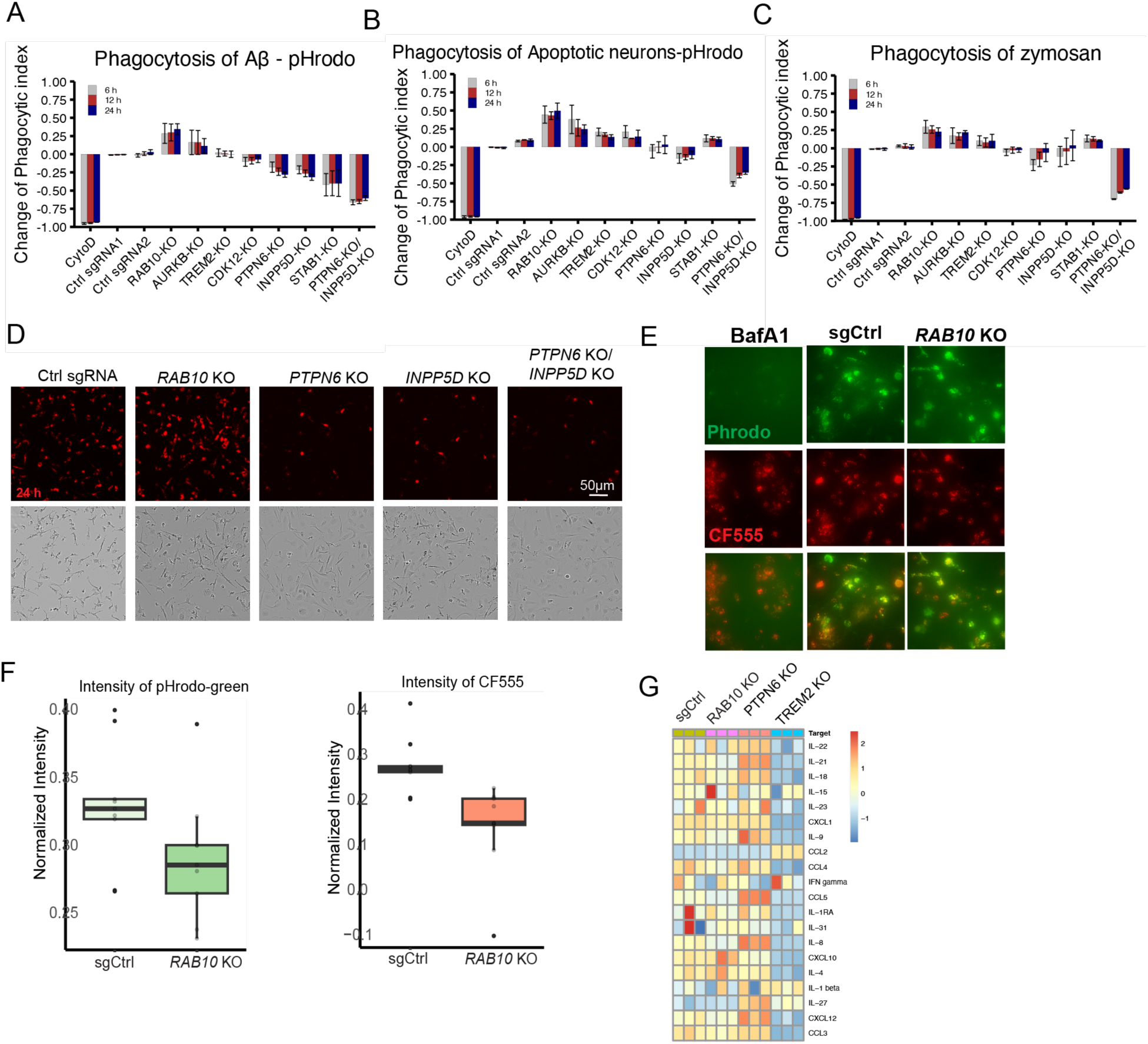
Targeted depletion of AD GWAS genes modulates microglial phagocytic capacity in a substrate-specific manner. **A-C.** Bar plots depict the phagocytic index of Aβ–pHrodo (A), apoptotic neurons (B) and zymosan particles (C) uptake at 6, 12, and 24 hours in iMicroglia for each gene knockout. Phagocytic index values were normalized to control sgRNA, with phagocytic index > 1 indicating increased cellular intensity and phagocytic index < 1 indicating reduced cellular intensity relative to control. Data are presented as mean ± SEM (n = 3). **D.** Representative phase-contrast and fluorescence images showing the Aβ-pHrodo accumulation 24 hours after Aβ-pHrodo addition in iMicroglia expressing control, *RAB10*, *PTPN6*, *INPP5D* or *PTPN6*/*INPP5D* sgRNAs. **E.** Representative IF images showing the lysosomal acidity in control sgRNA and *RAB*10 KO iMicroglia was determined using a ratiometric pH dyes (pHrodo Green dextran and CF555 dextran). Bafilomycin (BafA1, 1 µM, 3 hours) treated control iMicroglia served as a positive control. **F.** Boxplot showing the quantification of the intensity of pHrodo-green and CF555 from experiment in Fig. 6E. Both the intensity of pHrodo-green dextran and CF555-detran were decreased in *RAB10* KO cells, suggesting lysosomal deacidification and impairment of the endolysosomal pathway. The horizontal black lines within the boxes denote median values (50th percentiles), the black boxes contain the values from the 25th to the 75th percentiles and the black whiskers denote values at the 5th and 95th percentiles.The black circles represent the data distribution. **G.** Heatmap showing the release of cytokines and chemokines from iMicroglia expressing control, *RAB10*, *PTPN6* and *TREM2* sgRNAs (n=3).

*RAB10* depletion significantly increased phagocytic uptake across substrates, consistent with its known regulatory role in phagosome maturation and membrane recycling ^66^. Dual knockout of *INPP5D* and *PTPN6* produced a ∼60% reduction in Aβ uptake, substantially greater than the 20–30% reductions seen with either single-gene knockout (Fig. 6D, supplementary Fig. 7B), suggesting that SHIP1 (encoded by *INPP5D*) and SHP1 (encoded by *PTPN6*) may act cooperatively within inhibitory signaling pathways that regulate microglial phagocytosis ^67–69^.

Given the elevated lysosomal signatures in *RAB10*-deficient cells, we asked whether the increased Aβ-pHrodo signal reflected enhanced phagocytic uptake (potentially beneficial) or impaired cargo degradation (detrimental despite increased uptake) ^70^. A ratiometric assay using pH-sensitive (pHrodo-Green) and pH-insensitive (CF555) dextrans ^71^ revealed increased lysosomal acidification in *RAB10*-deficient cells despite reduced overall dextran uptake (Fig. 6E–F, Supplementary Fig. 7C), suggesting a broader disruption in endocytic trafficking rather than a simple increase in phagocytosis. Decreased EEA1 expression by IF further confirmed dysregulation of endosomal-lysosomal pathways in *RAB10*-depleted cells (Supplementary Fig. 7D), collectively indicating compromised endosomal-lysosomal functions.

In parallel experiments, *TREM2* depletion caused significant lysosomal dysfunction, evidenced by reduced LysoTracker Red signal relative to controls (Supplementary Fig. 7E–F). Consistent with this, *TREM2* knockout cells showed diminished fluorescence in a DQ™-BSA-red degradation assay, confirming impaired breakdown of internalized cargo (Supplementary Fig. 7G). Together, these results demonstrate that both *RAB10* and *TREM2* are critical for maintaining endolysosomal integrity, and that their depletion compromises substrate degradation and overall microglial phagocytic function. Finally, cytokine and chemokine release were slightly increased in *PTPN6* knockout cells and significantly decreased in *TREM2* knockout cells, indicating gene-specific effects on inflammatory output (Fig. 6G).

Together, these findings show that specific gene perturbations may drive coordinated changes in microglial morphology, transcriptional identity, and functional capacity. The observed relationships—loss of homeostatic gene expression, emergence of rounder cellular morphology, shifted inflammatory landscape, and reduced phagocytic activity— provide insight into how microglia transitions between functional states through concurrent shifts in morphology, inflammatory state, and transcriptional identity. This integrated view linking morphology, transcriptional state, and phagocytic performance highlights the inherent plasticity of microglia and their context-dependent roles in neurodegeneration and suggests that environmental or genetic perturbations can destabilize microglial homeostasis, dynamically shifting cells toward either more resilient and neuroprotective or more neurotoxic states in disease settings.

## Discussion

In this study, we established a streamlined and reproducible approach for generating mature iPSC-derived microglia that faithfully recapitulate key molecular, functional, and inflammatory features of human microglia, including their responses to Aβ, LPS, and Tau PFFs, providing a robust platform for modeling complex neurodegenerative disease processes. Our scRNA-seq data showed that iMicroglia exhibit a spectrum of distinct transcriptional states—homeostatic, antigen-presenting, cycling, lipid-associated, and DAM-like—mirroring the heterogeneity reported *in vivo* and in other iMicroglia systems ^45,46,52,72,73^. This diversity provides an essential foundation for interpreting how microglia respond to physiological and pathological cues. We further found that iMicroglia mount distinct, stimulus-specific transcriptional responses. Tau PFFs, an AD-relevant stimulus ^45,74^, induced broad inflammatory activation and depleted the states observed in untreated iMicroglia, whereas Tau monomers had minimal effect (Fig. 2). Tau PFF-treated cells separated from untreated clusters in UMAP space, reinforcing that iMicroglia do not collapse into a single reactive state upon stimulation ^45^. Notably, previous studies reported reduced antigen-presenting microglia (IBA1+/CD74-high cells in that report) and enriched DAM-like clusters under disease conditions ^53^, and we observed similar state shifts in our Tau PFF-treated iMicroglia. Prolonged Tau PFF exposure further enriched DAM and MGnD signatures, underscoring the relevance of this model for studying AD-associated microglial phenotypes. In contrast, fibrillar Aβ and apoptotic neurons elicited a more moderate shift toward an APOE/GPNMB-high (DAM-like) state, in agreement with previous findings ^45^. Together, these observations show that iMicroglia not only recapitulate human microglial state diversity but also undergo dynamic, stimulus-specific state transitions—plasticity that underscores the relevance of our platform for studying how microglia integrate environmental signals, shift between states, and contribute to neurodegenerative processes such as AD.

Microglial states arise from the intricate integration of epigenetic, transcriptomic, proteomic, and metabolic cues ^44^, and understanding how these transcriptionally defined states relate to microglial morphology and function—particularly under disease-relevant conditions—may offer a new angle for uncovering disease mechanisms and therapeutic opportunities. Our morphological profiling showed that GPNMB+ (DAM-like) iMicroglia exhibited larger, flatter, more amoeboid morphologies (low eccentricity, large area), while HLA+ (antigen-presenting) cells were typically smaller and more rounded (low eccentricity, small area). iMicroglia also underwent substantial morphological remodeling in response to stimuli: acute LPS or Tau PFF exposure triggered rod-like shapes and depleted IBA1⁺ homeostatic and HLA⁺ antigen-presenting cells, consistent with the corresponding transcriptomic changes. Similarly, *INPP5D* and *PTPN6* knockouts led to loss of both HLA-II+ and homeostatic populations, aligning with recent scRNA-seq studies reporting selective depletion of antigen-presenting microglial clusters in AD ^73^, and supporting earlier observations that HLA class II expression varies with physiological and pathological state and may serve as an indicator of homeostatic versus disease-associated microglial activation ^75^. By integrating morphological features with transcriptomic states, we were able to track global microglial state transitions and systematically analyze how microglia reorganize in response to diverse stimuli. Linking transcriptional states with morphological features in this way offers new insight into microglial plasticity and function across physiological and pathological settings; although this study focused on IBA1, GPNMB, and HLA, expanding to additional markers will further refine our understanding of microglial morphological and state transitions.

To examine the functional consequences of these morphological and state transitions further, we characterized selected AD risk gene knockout iMicroglia with altered phagocytic capacity. *RAB10* is a known regulator of endosomal trafficking across multiple brain cell types ^76,77^ and has been implicated in AD and PD ^78–80^, but its role in microglial phagocytosis has been poorly defined. *RAB10* plays a critical role in endoplasmic reticulum and endolysosomal function and has been linked to LRRK2-mediated manganese toxicity through dysregulation of the autophagy–lysosome pathway in microglia ^81–83^. Our findings provide new insight into the role of *RAB10* in regulating microglial function and highlight its potential as a therapeutic target. Depletion of *INPP5D* and *PTPN6*, both components of the ITIM signaling pathway, also impaired phagocytosis ^84^. *INPP5D*, a microglia-enriched gene associated with AD risk and amyloid pathology, shows context-dependent roles across studies ^64,69,85–87^. Consistent with previous findings ^86^, acute *INPP5D* depletion in iMicroglia significantly reduced Aβ uptake. *PTPN6*, another AD-associated gene and a potential *CD33* effector, similarly reduced phagocytosis, and combined *INPP5D/PTPN6* depletion exacerbated this effect, hinting on their coordinated action within a shared inhibitory signaling pathway ^84^. Further mechanistic studies will be important to elucidate how *INPP5D* and *PTPN6* jointly and independently regulate microglial function in neurodegeneration.

Given these functional changes, our morphological profiling revealed distinct changes in microglial plasticity. *PTPN6*-deficient iMicroglia displayed more of donut-shaped morphologies, increased GPNMB expression (a DAM-like state feature), and reduced Aβ uptake, suggesting a shift toward a less phagocytic state. In contrast, *RAB10*-deficient cells showed impaired phagocytic processing without major morphological changes. These observations support the growing view that microglial morphology and transcriptional states act in concert to shape functional capacity and may reflect specialized roles in pathology. Recent work has emphasized this concept of microglial morpho-functional plasticity, linking distinct morphological signatures to disease states and stimulus-specific responses ^88–90^ ^91^. Our findings reinforce this concept and highlight the value of high-content imaging-based morphological profiling for mapping microglial state dynamics and associated functions. More broadly, this work offers a framework for linking microglial morphology and states to disease-relevant behaviors and identifies molecular nodes that may be therapeutically targeted to restore or reprogram microglial function. As future studies expand to larger genetic perturbations, diverse environmental contexts, and additional state markers, this study will serve as a valuable resource for understanding how microglial plasticity contributes to both neuroprotection and neurodegeneration.

## Conclusions

Our study establishes a robust and versatile platform for linking microglial morphology, transcriptional states, and function. By integrating multi-omics profiling, morphological profiling, and targeted genetic perturbation, we show that microglial states are accompanied by coordinated morphological and functional changes, reflecting the morpho-functional plasticity observed *in vivo*. Mechanistic insights—including the shared inhibitory signaling mediated by *INPP5D* and *PTPN6*—highlight pathways that may underlie microglial dysfunction in AD. As this platform is expanded to additional genetic perturbations and stimuli, it will deepen our understanding of how microglial plasticity shapes resilience and vulnerability in neurodegenerative disease, and may guide strategies to reprogram microglial states for therapeutic benefit.

## Methods

### Human iPSC culture

In brief, human iPSC were maintained with daily feedings in TeSR™-E8™ media (05990; Stemcell technologies) on vitronectin (A14700; Thermo Fisher) coated (1 ug/cm^2^) tissue culture treated plastic. When iPSC cultures reached ∼75% confluency, they were subcultured by treating with ReLeSR™ (100-0483; Stemcell Technologies) for 5 minutes at room temperature. Cultures underwent a minimum of 3 passages after thawing before initiating differentiations.

### Differentiation and culture of iPSC-derived microglia

We adapted a pre-existing protocol to differentiate iPSCs into microglial precursors ^10^, and then further optimized the maturation protocol from microglial precursors to generate a more scalable and pure, homeostatic microglial monoculture. On day 0, ∼75% confluent iPSC cultures were dissociated into single cells using ACCUTASE™ (07920; Stemcell Technologies) and seeded at a density of 90,000 cells/cm^2^ on Geltrex™ (A1413201; Thermo Fisher) coated (60 µg/cm^2^) vessels using 0.2 mL/cm^2^ TeSR E8 media containing 60 ng/mL BMP4 (314-BP-010/CF; R&D Systems), 7.5 ng/mL Activin A (338-AC-010/CF; R&D Systems), 3 µM CHIR99021 (4423; Tocris), and 10 µM Y-27632 (S1049; Selleck Chemicals). After overnight incubation at 37°C with 5% CO_2_ for 18 hours, a complete media change was performed with 0.2 mL/cm^2^ TeSR™-E6 (05946; Stemcell technologies) media containing 80 ng/mL BMP4, 10 ng/mL Activin A, and 2 µM IWP2 (3533/10; R&D Systems). On day 2, a complete media change was performed with 0.2 mL/cm^2^ E6 media containing 80 ng/mL BMP4, 10 ng/mL Activin A, 2 µM IWP2, and 20 ng/mL bFGF (130-093-839; Miltenyi). On day 3 the cultures are dissociated into single cells using Accutase for 20 minutes followed by trituration with an equal volume of E6 media. After centrifugation (300g, 5 min.), the supernatant was aspirated and the pellet was resuspended and seeded in Geltrex coated vessels (60 µg/cm^2^) at 55,000 cells/cm^2^ using 0.2 mL/cm^2^ E6 media containing 15 ng/mL VEGF (293-VE-010/CF; R&D Systems), 10 µM Y-27632, and 5 ng/mL bFGF, or cryopreserved in E6 media containing 15 ng/mL VEGF, 10 µM Y-27632, 5 ng/mL bFGF and 8.5% DMSO. On day 4, a complete media change was performed with 0.2 mL/cm^2^ E6 media containing 15 ng/mL VEGF, and 5 ng/mL bFGF. On days 5 and 6, a complete media change was performed with 0.2 mL/cm^2^ E6 media containing 15 ng/mL VEGF, 5 ng/mL bFGF, 200 ng/mL SCF (255-SC-200/CF; R&D Systems), and 20 ng/mL IL-6 (206-IL-050/CF; R&D Systems). On day 7, a complete media change was performed with 0.2 mL/cm^2^ E6 media containing 100 ng/mL SCF, 10 ng/mL IL-6, 30 ng/mL TPO (288-TP-025/CF; R&D Systems) and 30 ng/mL IL-3 (203-IL-010/CF; R&D Systems) and returned to the incubator for 2 days. On day 9, suspended cells were collected from the supernatant by centrifugation at 300g for 5 min. A complete media change was performed with 0.2 mL/cm^2^ E6 media containing 100 ng/mL SCF, 10 ng/mL IL-6, 30 ng/mL TPO and 30 ng/mL IL-3, and the suspended cells were returned to the fresh media in the culture vessels. From days 10-14 the volume was increased by 25% each day with the addition of fresh E6 media using the total volume/vessel to achieve final concentrations of 100 ng/mL SCF, 10 ng/mL IL-6, 30 ng/mL TPO and 30 ng/mL IL-3. On day 14 the cells in suspension were harvested and centrifuged at 300g for 5 min. (adherent cells were then discarded). Pellets were then re-suspended in R-10 media (RPMI +glutamax containing heat inactivated 10% FBS (A4766801; Life Technologies), 100 ng/mL IL-34 (5265-IL-010/CF; R&D Systems) and 10 ng/mL M-CSF (216-MC-025/CF; R&D Systems)) and seeded into fibronectin (33016015; Gibco) coated (2 µg/cm^2^ in PBS at 37°C for 3 hrs followed by 1 wash with TC grade water) vessels at a seeding density of 40,000 cells/cm^2^ in a volume of 0.2 mL/cm^2^. On day 16 and 18, to maintain the loosely adherent and suspended cells, 0.1 mL/cm^2^ R-10 media was added directly, replenishing 100 ng/mL IL-34 and 10 ng/mL M-CSF for the full volume. On day 20, the supernatant containing non-adherent cells was collected. 0.1 mL/cm^2^ cold PBS was then added (10 min. at 4°C) to the adherent cells, allowing them to be easily removed by pipetting. The adherent and non-adherent cells were pooled and either re-plated at 75,000 cells/cm^2^ on fibronectin coated vessels using R-6 media (RPMI +glutamax containing 6% FBS, 100 ng/mL IL-34, 10 ng/mL M-CSF and 25 ng/mL TGF-β1 (240-B-010/CF; R&D Systems)) or cryopreserved (R-10 media with 8.5% DMSO). On day 22 a half media change was performed using R-2 media (RPMI with glutamax containing 2% FBS, 100 ng/mL IL-34, 10 ng/mL M-CSF and 50 ng/mL TGF-β1). On day 24 a half media change was performed using a serum free (SF) media formulation consisting of DMEM/F-12, N2, GlutaMAX™ (35050061; Gibco), and 2-Mercaptoethanol (21985023; Gibco) supplemented with 200 ng/mL IL-34, 20 ng/mL M-CSF and 100 ng/mL TGF-β1. On day 26, a complete media change was performed using SF media supplemented with 100 ng/mL IL-34, 10 ng/mL M-CSF and 50 ng/mL TGF-β1. Half media changes were performed every other day using SF media supplemented with 200 ng/mL IL-34, 20 ng/mL M-CSF and 100 ng/mL TGF-β1.

### Human monocyte–derived macrophage culture

Isolation and culture of human monocyte–derived macrophages (hMDMs) were performed following the previous protocol ^92^. Basically, human peripheral blood mononuclear cells (PBMCs) were isolated from consenting healthy donor buffy coats using Ficoll Paque Plus (Sigma-Aldrich). Next, CD14+C16− monocytes were isolated from PBMCs using the classical monocyte isolation kit (Miltenyi Biotec) per kit instructions. Purified monocytes were collected and centrifuged at 300 × g for 10 min and then resuspended in the growth medium. Cells were diluted to a final concentration of 10% DMSO, frozen, and stored in liquid nitrogen for future use. For each experiment, aliquots from mul-tiple donors were thawed, and human monocyte–derived macrophages (hMDMs) were differentiated in culture for 5–7 d in the growth medium high-glucose DMEM containing 10% heat-inactivated serum and 50 ng/ml recombinant human M-CSF (PeproTech) with partial media changes every other day.

### Immunocytochemistry

Cells were fixed with fresh 4% formaldehyde for 15 minutes at room temperature. Fixed cells were washed three times with PBS for 10 minutes and incubated with SuperBlock™ Blocking Buffer supplemented with 0.1% Triton X-100 for 30 minutes at room temperature. Samples were then incubated with primary antibodies at 4℃ overnight, washed three times with PBS for 15 minutes at room temperature and incubated with secondary antibodies for our hour at room temperature. Lastly, samples were incubated with 1 µg/ml DAPI for 10 minutes at room temperature and washed three times with PBS for 10 minutes. Samples were imaged using Zeiss LSM980 confocal microscope.

### pHrodo-Red labeling

Fibrillar Aβ was generated from Beta-Amyloid (1-42) (Anaspec, AS-20276) following the protocol^93^. Briefly, 0.25 mg of Beta-Amyloid was reconstituted in 40 µl of DMSO and 460 µl of PBS for 24 hours at 37℃ on a rotator. Reconstituted Aβ solution was then centrifuged at 16,000 g for three minutes to collect the aggregates. Supernatant was carefully aspirated to avoid disturbing the pellet. The pellet was then rinsed with 1 ml of HBSS for three times and resuspended in 200 µl of 0.1 M sterile sodium bicarbonate and mixed well. 6 µl of pHrodo Red dye (10 nM, Invitrogen, P36600) was added to the Aβ aggregate solution. The reaction tube was incubated on a rotator at room temperature in the dark for 15 minutes. The reaction solution was then centrifuged at 16, 000 g for 1 minute and pHrodo-labeled Aβ will aggregate into a dark purple pellet. The supernatant was aspirated carefully and the Aβ pellet was washed four times with 1 ml HBSS at room temperature. After the last wash, the pellet was resuspended with 550 µl of HBSS and aliquoted into EP tubes and saved in -80℃ until being used.

### Phagocytosis assay

Differentiated microglia at Day 20 were harvested and re-plated in 96-well tissue culture plates at a density of 30,000 cells per well. Cells were cultured as described above for 10 days. At day 30, LPS (100 ng/ml) was added to selected wells to pre-treat the cells. At day 31, the phagocytosis substrates A**β**-pHrodo (1µM), zymosan (3.75 µg/mL) or apoptotic neurons-pHrodo (0.44 mg/mL) were added the cultures and transferred to the Incucyte automated microscope and imaging intervals were set at every 2 hours. The first time point (*t* = 0) was acquired within 20 min of the addition of the substrates. Total Red Object Integrated Intensity (RCU x µm²/Image) of a well was calculated by the automatic metric over five fields per well. At least three wells were used per condition. Images were acquired at 10X objective at 800 ms (red) exposures. After the last time point, nuclear green LCS1 (1:750, ab138904) was added to the cells and incubated for 15 minutes before the last image acquisition to determine total cell number per field. To calculate a phagocytic index for each condition, the total pHrodo-positive intensity per well over five fields was determined by the automatic metric of the Incucyte software and normalized to the total cell number per well over the same five fields. The phagocytic index was then calculated by normalizing each group of the experiment to the control group. Experiments were performed for three biological replicates using three independent differentiations.

### Human cytokine and chemokine assay

iMicroglia was cultured in 96 well plates and fed with fresh media. Experimental groups were incubated with LPS (100 ng/ml), fibrillar tau (10 µg/ml) or monomeric tau (10 µg/ml) for 24 hours. Non-treatment (control) groups were added the same amount of PBS. 100 µl of the conditioned media was collected from each group and analyzed using the Millipore Premixed 30 Plex kit following the manufacturer’s protocol by Flow Core of Genentech. At least three replicates were processed for each group. Experiments were performed for three biological replicates using three independent maturations.

### Morphological profiling of iMicroglia

Cell segmentation models were trained via cellpose2 with a foundation of the cyto model. Almost all cells were IBA1+, and can be segmented with the IBA1 model. GPNMB+ and HLA+ cells were also segmented via cellpose models, and parent-children relationships were assigned to IBA1-GPNMB or IBA1-HLA pairs to determine IBA1+GPNMB-HLA-, GPNMB+HLA+ populations. After cell segmentation, features related texture, radial distribution, intensity, AreaShape were extracted from all four channels (DAPI, GPNMB, HLA, and IBA1) via CellProfiler 4. Well-level aggregation was performed via signed KS normalization^94^. To identify features that differed the most upon treatments, one-way ANOVA was performed between treatment group and untreated group followed by post hoc Tukey analysis. To address multiple comparisons, adjusted p values were calculated via the Benjamini-Hochberg procedure. For single cell level analysis, outliers (below 5 percentile and above 95 percentile) were removed, and all features were transformed to Z scores. For PHATE dimension reduction, features with a correlation value larger than 0.9 were iteratively removed, and PHATE was run with the default parameters.

### CRISPR ribonucleoprotein (RNP) method

Day 20 iMicroglia were thawed and seeded at 75,000 cells/cm^2^ in R-6 media on fibronectin coated vessels. On day 22 a half media change was performed using R-2 media. On day 24, after 4 days of recovery from thaw, supernatants containing cells in suspension were collected. Remaining adherent cells were treated with cold PBS at 4°C for 10 minutes (to enhance physical retraction from the plate via non-enzymatic methods) and dissociated by pipetting. The dissociated adherent cells were combined with the suspension fraction and placed on ice. 5 µL nucleofection reaction mixtures were prepared for each well of a 16-well Nucleocuvette Strip (Lonza) using 3 µL of 100 µM guide RNA (Synthego), 1 µL of 10 nM Alt-R® Cas9 nucleofection Enhancer (1075916; IDT), 1 µL of 10 µg/mL Alt-R™ S.p. Cas9 Nuclease V3 (1081059; IDT) and incubated at room temperature for 20 minutes. During this time, iMicroglia cells prepared for nucleofection were centrifuged at 300g for 5 minutes. The supernatant was aspirated and the pellet was resuspended in P3 Primary Cell Nucleofector Solution (Lonza) at 300,000 cells/15.2 µL. 15.2 µL of cells in the P3 buffer were then added to each 5 µL reaction mix before adding to each well of a nucleofection cassette. The cassette was inserted into the 4D-Nucleofector X Unit (Lonza) and pulsed on program EM 110. The cassette wells were then gently flooded with 100 µL R-2 media. Electroporated cells were transferred to microcentrifuge tubes for post nucleofection cell counts. Cells were then suspended in half R-2/half SF media containing 100 ng/mL IL-34, 10 ng/mL M-CSF and 50 ng/mL TGF-β1 and plated into 96 well plates (6055302; Revvity) at a density of 90,000 cells/cm^2^. On day 26 (2 days post nucleofection), a complete media change was performed using SF media supplemented with 100 ng/mL IL-34, 10 ng/mL M-CSF and 50 ng/mL TGF-β1. Half media changes were performed every other day using SF media supplemented with 200 ng/mL IL-34, 20 ng/mL M-CSF and 100 ng/mL TGF-β1. Cells are ready for phagocytosis assay by day 29.

### Bulk RNAseq sample preparation

RNA was purified from microglia using the Zymo Directed RNA miniprep kit (R2050; Zymo Research) following manufacturer’s instructions. Briefly, microglia were cultured in 96 well plates until the day of harvest. To harvest the samples, culture media was removed from the wells and 100 µl of TRIi reagent was added to lyse the cells. Cell lysis can either be frozen at -80℃ or proceed to RNA purification following the Zymo directed protocol. RNA concentration was determined by Nanodrop. At least 100 ng of purified RNA was submitted for the RNAseq. At least three biological replicates were performed for each sample. Sequencing was performed by the NGS core of Genentech.

### Single cell RNA-seq sample preparation

iMicroglia were cultured in 24 or 12 well plates until the desired time point. Treatment groups were incubated with stressors such as LPS (20 ng/ml), Tau PFF (10 µg/ml) or monomeric tau (10 µg/ml) for 24 hours before the time of harvest. 100,000 to 200,000 iMicroglia were harvested with cold PBS and transferred to 1.5 ml EP tubes. iMicroglia were centrifuged at 400 g for 7 minutes and washed with cold PBS supplemented with 1% BSA for three times. After the last wash, iMicroglia pellets of each sample were resuspended in 100 µl Staining buffer (Biolegend, 420201). 0.5 µg (1 µl) of a unique Cell Hashing antibody (TotalSeq™-A Antibodies, Biolegend) and 1 µM of LIVE/DEAD™ Fixable Far-Red Dead Cell Stain Kit (Invitrogen, L10120) were added to each sample and mixed well. Samples were incubated at 4℃ for 20 minutes and then washed for three times with 1 ml Staining buffer. After the last wash, cells were resuspended in 200 µl PBS with 1% BSA and proceeded with FACS sorting using the far red channel. 20, 000 Live cells for each sample were sorted and kept in individual EP tubes. Four samples were pooled for one standard 10X run. 49,560 viable cells (aiming for recovery of 30,000 cells for each run) were processed using the Chromium Next GEM Single Cell 3’ Reagent Kits v3.1 to generate cDNA libraries following manufacturer’s instructions. The quality and quantity of cDNA and libraries were evaluated by Tapestation.

### RT-qPCR

RNA was purified using the RNeasy Micro Kit (74004; Qiagen) following the manufacturer’s protocol. qScript XLT 1-Step RT-qPCR ToughMix (95133-500; Quantabio) was used to run the reverse transcription quantitative PCR. 10 ng of purified RNA was used per reaction. Two technical replicates and three biological replicates were run for each assay. The assays were run in a ViiA 7 Real-Time PCR System using Tagman primers with the parameters: the parameters: cDNA synthesis, 50°C for 10 minutes, initial denaturation, 95°C for 1 minutes, PCR cycling (40 cycles), 95°C for 10 seconds, 60°C for 30s.

### Lysosomal luminal pH assessment

iMicroglia cultures were treated with both 10 µg/mL pHrodo™ Green Dextran (Invitrogen; P35368) and 10 µg/mL CF®555 (Biotium; 80112) dyes for 18 hours. The cultures were then washed with fresh culture media and 100 nM Bafilomycin (Cell Signaling; 54645S) was added to control wells. 3 hours later, the cultures were imaged live (Leica Thunder) prior to fixation (15 minutes in 4% Paraformaldehyde).

### Lysosomal hydrolytic activity assay

iMicroglia cultures were treated with both 25 µg/mL DQ™ Red BSA fluorogenic substrate (Invitrogen; D12051) and 25 µg/mL Albumin from Bovine Serum (BSA) Alexa Fluor™ 488 conjugate (Invitrogen; A13100) for 1.5 hours. Control cultures were additionally treated with 100 µM Leupeptin Protease Inhibitor (Thermo Scientific; 78435) and 20 mM NH_4_Cl (Sigma; A9434). Live imaging on the Leica Thunder was performed following two PBS washes along with the addition of fresh culture media.

### Lysosomal acidity assessment

iMicroglia cultures were treated with either 100 nM LysoTracker™ Red DND-99 (Invitrogen; L7528) or 100 nM LysoSensor™ Green DND-189 (Invitrogen; L7535) for 30 minutes. Control wells were pre-treated with 100 nM Bafilomycin. Cells were imaged live on Leica Thunder. Experiments were performed for two biological replicates using cells from two differentiations.

### Tri-culture

iMicroglia were cultured up to day 24 for tri-culture using methods described in this paper. Astrocyte precursor cells (APCs) were derived based on methods described in Russo et al ^95^. Briefly, iPSCs were differentiated into NPCs using the Stemdiff SMADi Neural Induction kit (StemCell Technologies; Cat.No. 08581) according to the manufacturer’s instruction. Neural rosettes were manually picked and expanded in NPC media (DMEM/F12, Neuralbasal, Glutamax, B27, N2) supplemented with FGF2, EGF and BDNF until confluency. NPCs were then dissociated as single cells, and seeded into spheroids using the AggreWell 800 plates (StemCell Technologies; Cat. No. 34815) at a density of 10,000 cells per microwell. Spheroids were initially grown in the NPC media for one week. In the subsequent two weeks, media were slowly switched to the astrocyte growth media (Lonza; Cat. No. CC-3186) with three quarters media change. After three weeks on AggreWell, spheroids were transferred onto Geltrex coated plates at 1:6 split ratio and allowed continued growth without re-plating for 4 weeks in Astrocyte growth media (Lonza; Cat. No. CC-3186). By the end of week 4, cells are differentiated into astrocyte precursor cells based on astrocyte lineage marker expressions such as GFAP, S100B and AQP4. APCs were expanded for two more passages and dissociated into single cells for cryopreservation. Cells were fed three times a week at all stages except for before the NPC stage. iNGN2 neurons were generated using the protocol described in our previous publication ^96^. We generated iPSC derived excitatory neurons (iNGN2) and cultured them at 120,000 cells/cm2 to day 14 before addition of iAstrocytes.

Tri-culture: At day 14 of iNGN2 cultures, cryopreserved astrocyte progenitor cells (APC) were thawed and directly seeded into iNGN2 cultures at 15,000 cells / cm2, concurrent with a half media exchange using tri-culture media (DMEM/F12, Glutamax, BME, N2) supplemented with 20 ng/mL BDNF and 20 ng/mL GDNF. Additional half media exchanges using tri-culture media with ligands were performed on days 16, 18, and 20. iMicroglia were directly seeded into iNGN2/iAstrocytes co-cultures at 120,000 cells / cm2 in tri-culture media containing IL-34, M-CSF and TGF-β1. Half volume media exchanges were performed on tri-cultures every other day using tri-culture media containing IL-34, M-CSF and TGF-β1 until day 28.

### Bulk RNA-seq analysis

RNA-sequencing data were analyzed using HTSeqGenie (Pau and Reeder, 2012) in Bioconductor as follows: first, reads with low nucleotide qualities (70% of bases with quality <23) or matches to rRNA and adapter sequences were removed. The remaining reads were aligned to the human reference genome (human: GRCh38.p10) using GSNAP (PMID:20147302, 27008021) version ‘2013-10-10-v2’, allowing maximum of two mismatches per 75 base sequence (parameters: ‘-M 2 -n 10 -B 2 -i 1 -N 1 -w 200000 -E 1 --pairmax-rna=200000 --clip-overlap’). Transcript annotation was based on the Gencode genes database (human: GENCODE 27). To quantify gene expression levels, the number of reads mapping unambiguously to the exons of each gene was calculated.

For expression level plots nRPKM values were computed as described in Law et al 2014 ^97^ and plotted in log2 scale. For heatmap representation the values were z scaled. Differential expression was performed with voom + limma as described in Law et al 2014. GO enrichment analysis was performed using clusterProfiler (V4.10 Bioconductor) package using all GO annotations from org.Hs.eg.db package (V3.18.0 Bioconductor).

For comparison to human microglia, we utilized multiple human in vivo datasets Herring et. al.^98^, Ramos et. al. ^99^, Trivino et. al. ^100^, Velmeshev et. al. ^101^, Bhaduri et. al. ^102^. A highly variable gene set was obtained using the pseudobulked microglia and astrocytes from each donor. Spearman correlation value of the highly variable geneset was used as a similarity measure and the correlation between microglia samples across donors and microglia vs astrocyte samples were used as positive and negative controls.

Brain vs peripheral myeloid cell data ^103^ was used to obtain a marker gene list for brain specific myeloid cell population as well as yolk sac specific population and two types of peripheral myeloid cell using the annotations provided by the authors. Gene set scores were calculated by taking the mean expression of the genes in each set. All bulk RNA-seq data were acquired from three biological replicates per line.

### Single cell RNA-Seq analysis

Sequencing data were analyzed using the Cell Ranger software suite (version 7.1.0). The workflow included the following steps: Raw FASTQ reads were aligned to the human reference genome GRCh38. The Gene expression was quantified using Cell Ranger’s default settings, excluding introns.

For downstream processing of the count data with scran(V1.30.2, Bioconductor), scater (V1.30.1, Bioconductor) and Seurat(V5.0.1) using the default settings of the package. For cell level QC we performed clustering with the filtered output from Cell Ranger using scran workflow. We then removed clusters that had high mitochondria % or low counts compared to other clusters. The data were then further filtered using mitochondrial read < 20% and detected genes > 1000. Filtered data were re-analyzed for final cluster annotations.

When combining data across multiple sequencing runs, the batch effect normalization was performed using Harmony^104^ as implemented in the R package (V0.1). For comparing cell states across datasets, metaneighbor (V1.22 Bioconductor)^105^ was used which identifies cell types with high similarity. Metaneighbor was run with highly variable genes, calculated with variableGenes function in the Metaneighbor package, as input.

For calculating gene set scores for single cell datasets, UCell was used (UCell V2.6.2, Bioconductor)^106^ using the top 50 markers provided by the original authors of Sun et. al.^52^.

All data were acquired from three biological replicates per line.

### Proteomics

#### Proteomics Sample Preparation

Cell pellets were lysed in denaturing buffer containing 8M urea, 150mM NaCl, 50mM HEPES pH 8.0), complete-mini (EDTA free) protease inhibitor (ROCHE), phosphatase inhibitor (PHOSstop, ROCHE), and benzonase nuclease (Sigma-Aldrich). Lysates were high-speed centrifugated (18,000 x g, 15 min) to remove insoluble material and protein concentrations were then determined by bicinchoninic acid assay. 60 μg of protein lysate per sample, save for samples NT-s2-1 and NT-s2-2 where all available material 16 μg and 4 μg, respectively, was subjected to cysteine disulfide reduction (4.1 mM dithiothreitol, 60 min at 37°C) and alkylation (9.1 mM iodoacetamide, 15 min at room temperature), and then diluted 4-fold with 50mM HEPES prior to enzymatic digestion. Overnight digestion was performed at 37°C using a combination of lysyl-endopeptidase (Wako) and sequencing grade trypsin (Promega), both at an enzyme to protein ratio of 1:50, the latter of which was added to the sample 3 h post incubation with the former. Resultant peptides were acidified with 20% TFA to a final concentration of 1%, clarified via high-speed centrifugation (18,000 x g, 10 min) prior to desalting via solid phase extraction (SOLA HRP Thermo). Desalted peptides were subsequently lyophilized to dryness^107^.

Each sample was resuspended with 100 µL of 100 mM HEPES pH 8.0 and labeled with a full vial of TMTPro 18-plex labeling reagent (Thermo) for 1 hr. 17 of 18 tags were utilized (TMT-126 to TMT-134C) corresponding to 17 samples. The reaction was quenched by the addition of 5 µL of 5% hydroxylamine (15 min at room temperature), combined and desalted by C18 Sep-Pak (Waters) solid phase extraction and eluents were dried to completion.

The TMT-labeled sample was redissolved in 0.15% trifluoracetic acid and subjected to centrifugation at 16,000 x g to remove insoluble material with the supernatant taken forward for offline high pH reversed phase liquid chromatography separation on an Agilent 1100 series HPLC system. Peptides were loaded onto an Agilent Zorbax 300 Extend C-18 analytical column (2.1 × 150 mm and 3.5 μm particle size, Part No: 763750-902). Solvent A consisted of 25 mM Ammonium Formate (pH 9.7), while solvent B was 100% acetonitrile. Peptide fractions were collected in 45 s intervals for a total of 63 min with a linear gradient of 15–60% solvent B. In total 96 fractions were collected and every 25th fraction was combined to form a final set of 24 fractions. Samples were dried in a speedvac, desalted using C18 StageTips as previously described and injected for nLC-MS/MS analysis.1 All data were acquired from three biological replicates per line.

### Proteomic Mass Spectrometry Analysis

nLC-MS/MS analysis was performed for each fraction on an Orbitrap Eclipse mass spectrometer (ThermoFisher) coupled to a Dionex Ultimate 3000 RSLC nano Proflow system (ThermoFisher) equipped with an Aurora Series 25 cm x 75 um I.D. column (IonOpticks). Low pH reversed-phase separation was performed at 300 nL/min on a 73.9 min two step linear gradient where solvent B (0.1% FA/98%ACN/2% water) was ramped from 4% to 30% over 68 min and then from 30% to 75% over 5.9 min with a total run time of 95 min. For these analyses, a real-time search against a human database was employed using an in-house instrument API program called InSeqAPI.2,3 Protein-closeout was implemented where proteins were placed on closeout list following detection of 3 distinct peptides, 5 unique peptides, and 10 peptide spectral matches. Intact peptides were surveyed in the Orbitrap (250% normalized AGC target, 120,000 resolution) and the top 10 peptides were selected for MS2 fragmentation (150% normalized AGC target; CID, 30 NCE) and analyzed in the ion trap. Synchronous-precursor-selection (SPS) MS3 scans were analyzed in the Orbitrap at 50,000 resolution with the top 8 most intense ions in the MS2 spectrum subjected to HCD fragmentation at a normalized collision energy of 45%, an AGC target of 3.0 x 105, and a max injection time of 400 ms^108–110^.

### Proteomic Data Analysis

MS/MS spectra were searched using COMET against a concatenated target-decoy database of human proteins (UniProt downloaded January, 2023) containing common contaminant sequences. Search parameters included trypsin cleavage with an allowance of 1 missed cleavage event, a peptide mass tolerance of 20 ppm, and recommended fragment ion settings for low resolution MS/MS with a fragment ion bin tolerance and bin offset of 1.0005 and 0.4, respectively. Searches permitted variable modifications of methionine oxidation (+15.9949 Da) and TMT labeled tyrosine (+304.20715 Da) with static modifications for cysteine carbamidomethylation (+57.0215 Da) and TMT tags on lysine and peptide N termini (+304.20715 Da). A false discovery rate (FDR) of 2% was applied at the peptide level to filter peptide spectra matches (PSMs). TMT reporter ions produced by the TMT tags were quantified with an in-house software package known as Mojave by calculating the highest peak within 20 ppm of theoretical reporter mass windows and correcting for isotope purities.4

The R package MSstatsTMT v.2.0.1 was used to preprocess PSM-level quantification before statistical analysis, to have protein quantification and to perform differential abundance analysis.5 MSstatsTMT estimated log2(fold change) and the standard error by linear mixed effect model for each protein. The inference procedure was adjusted by applying an empirical Bayes shrinkage. To test two-sided null hypothesis of no changes in abundance, the model-based test statistics were compared to the Student t-test distribution with the degrees of freedom appropriate for each protein and each dataset. The resulting P values were adjusted to control the FDR with the method by Benjamini– Hochberg.

## Key Resources Table

**Table 1.** Antibodies.

| Antibody | Dilution | Source | Identifier |
| --- | --- | --- | --- |
| IBA1 | 1:1000 | Synaptic systems | 234308 |
| GPNMB | 1:200 | R&D systems | AF2550 |
| HLA-DRA | 1:200 | Abcam | EPR3692 |
| P2RY12 | 1:200 | Novus Biologicals | NBP2-33870 |
| TMEM119 | 1:200 | Biolend | 853302 |
| RAB10 | 1:200 | Cell signaling | 8127S |
| SHIP1 | 1:200 | Cell signaling | 2727S |
| EEA1 | 1:100 | BD Biosciences | 610456 |

**Table 2.** Reagents for cell culture.

| Reagents | Source | Identifier |
| --- | --- | --- |
| LPS, E. Coli O111:B4 | EMD Millipore | LPS25 |
| cytochalasin D | Tocris Bioscience | 1233/1 |
| Monomeric Tau | SressMarq | SPR-479 |
| Recombinant fibrillar Tau | SressMarq | SPR-480 |
| Beta-Amyloid (1-42) | Anaspec | AS-20276 |
| BMP4 | R&DSystems | 314-BP-010/CF |
| Activin-A | R&DSystems | 338-AC-010/CF |
| CHIR 99021 | Tocris | 4423 |
| ROCKi (Y-27632) | STEMCELLTechnologies | 72302 |
| IWP-2 | R&DSystems | 3533/10 |
| FGF2 | Mylteni | 130-093-839 |
| VEGF | R&DSystems | 293-VE-010/CF |
| SCF | R&DSystems | 255-SC-200/CF |
| IL-6 | R&DSystems | 206-IL-050/CF |
| TPO | R&DSystems | 288-TP-025/CF |
| IL-3 | R&DSystems | 203-IL-010/CF |
| IL-34 | R&DSystems | 5265-IL-010/CF |
| M-CSF | R&DSystems | 216-MC-025/CF |
| TGFB-1 | R&DSystems | 240-B-010/CF |
| TeSR-E8 | STEMCELLTechnologies | 5990 |
| E8 | ThermoFisher | A1517001 |
| TeSR-E6 | STEMCELLTechnologies | 5946 |
| E6 | ThermoFisher | A1516401 |
| RPMI1640 with glutamax | ThermoFisher | 61870-127 |
| Matrigel | Gibco | 354277 |
| Fibronectin | Gibco | 33016015 |
| Vitronectin | Gibco | A14700 |
| Geltrex | Gibco | A14133-01 |
| FBS | Life Technologies | A4766801 |
| DMEM/F12 | Gibco | 21331-020 |
| Glutamax | Gibco | 35050-061 |
| BME | Gibco | 21985-023 |
| N2 | Gibco | 17502-048 |
| pHrodo® Red Zymosan<br>Bioparticles® | Sartorius | 4617 |
| DMEM/F12(no glutamine) | LifeTechnologies | 21331020 |
| B27without VA | Gibco | 12587010 |
| N2Max | R&DSsystems | AR009 |
| GlutaMAX | Gibco | 35050061 |
| NEAA | Gibco | 11140076 |
| SB431542 | STEMCELLTechnologies | 72232 |
| XAV939 | STEMCELLTechnologies | 72672 |
| DAPT | STEMCELLTechnologies | 72082 |
| Noggin | Mytenyi | 130-108-982 |
| Doxycycline | R&DSsystems | 4090/50 |
| DPBS (without Ca<sup>2+</sup>+Mg<sup>2+</sup>) | Gibco | 14190144 |
| iMatrix-511 | Takara | 892012 |
| Formaldehyde, 16% | Pierce | 28908 |
| ReLeSR | STEMCELL Technologies | 5872 |
| Arabinocytidine | Sigma | C1768 |
| dcAMP | Sigma | D0627 |
| Ascorbic Acid | Sigma | A4403 |

**Table 3.** Reagents for assays.

| **Reagents** | **Source** | **Identifier** |
| --- | --- | --- |
| LysoTracker™ Deep Red | Thermo Fisher | L12492 |
| Bafilomycin A1 | Cell signaling | 54645S |
| DAPI | 1:2000 | Invitrogen |
| Nuclear Green LCS1 | 1:700 | Abcam |
| Tagman probe | Thermo Fisher | 4453320 |
| Chromium Next GEM Automated<br>Single Cell 3' Library Kit v3.1 | 10X Genomics | 1000138 |
| Phrodo Green Dextran | Thermo Fisher | P35368 |
| DQ™ Red BSA | Thermo Fisher | D12051 |
| sgRNA | Synthego |  |
| Alt-R S.p. Cas9 Nuclease V3 | IDT | 1081059 |
| Alt-R® Cas9 Electroporation<br>Enhancer | IDT | 1075916 |
| qScript™ XLT One-Step RT-qPCR<br>ToughMix®, Low ROX™ | QuantaBio | 95134-500 |

**Table 4.** Cell lines.

| Cell line | Source | Identifier |
| --- | --- | --- |
| iPS11 | ALSTEM | iPS11 |
| KOLF2.1J | Jackson Laboratory | RRID: CVCL_B5P3 |
| iPS11N | ALSTEM | iPS11N |
| iPS26 | ALSTEM | iPS26 |

**Table 5.**
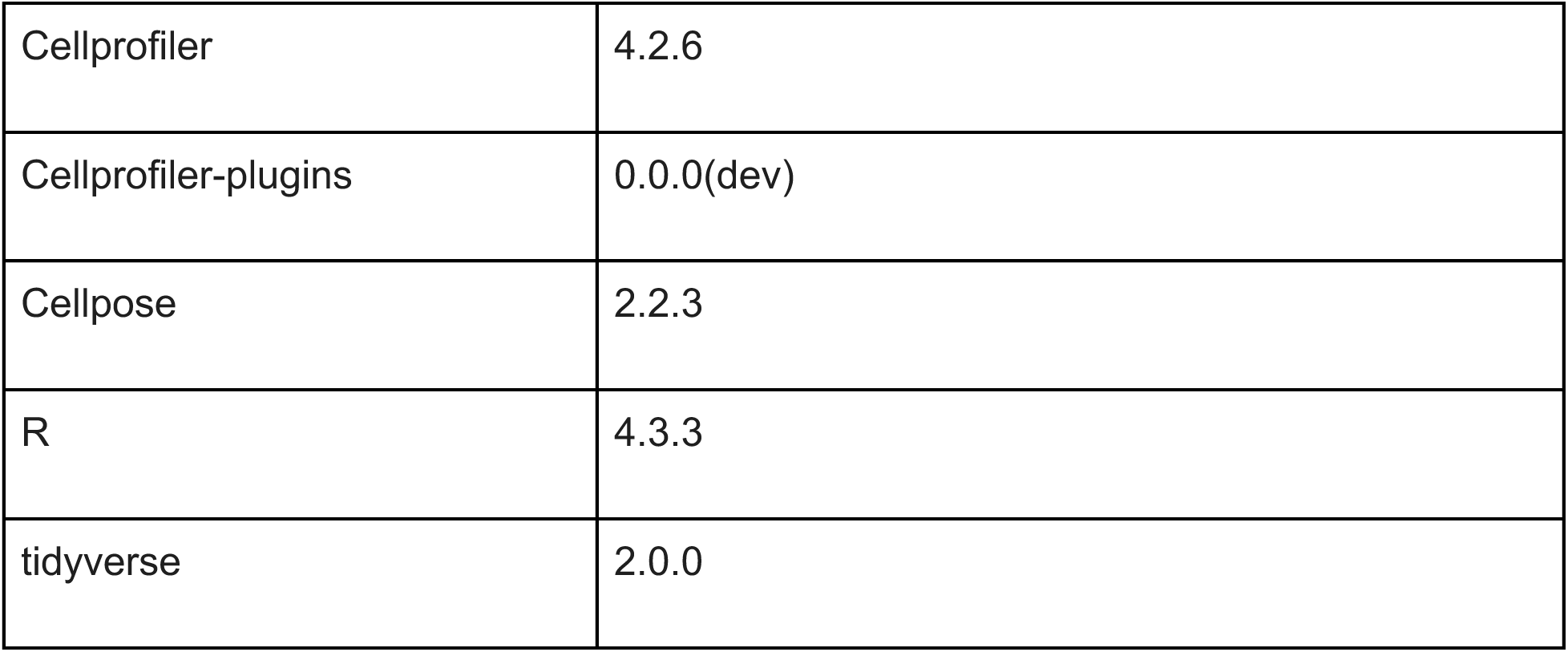
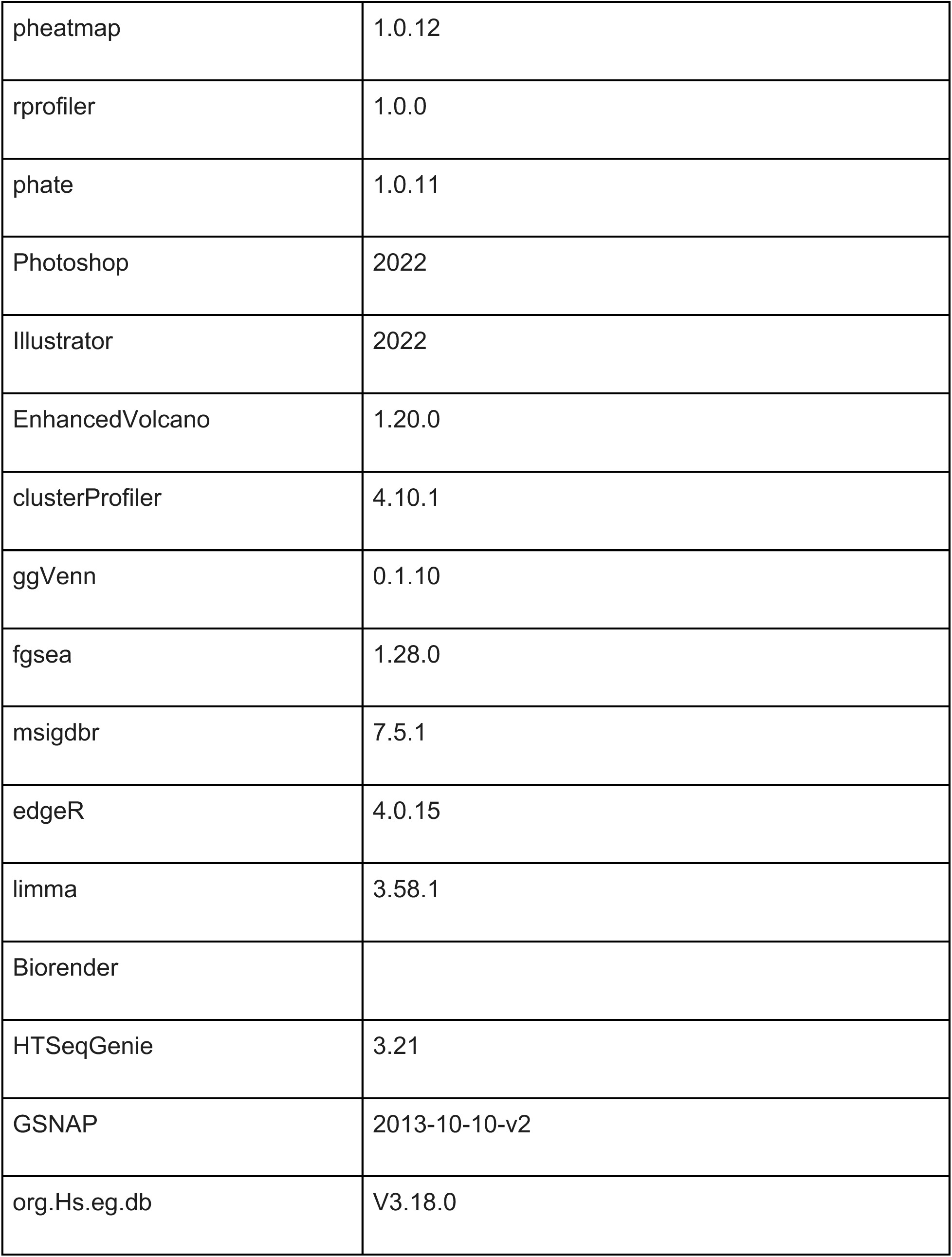
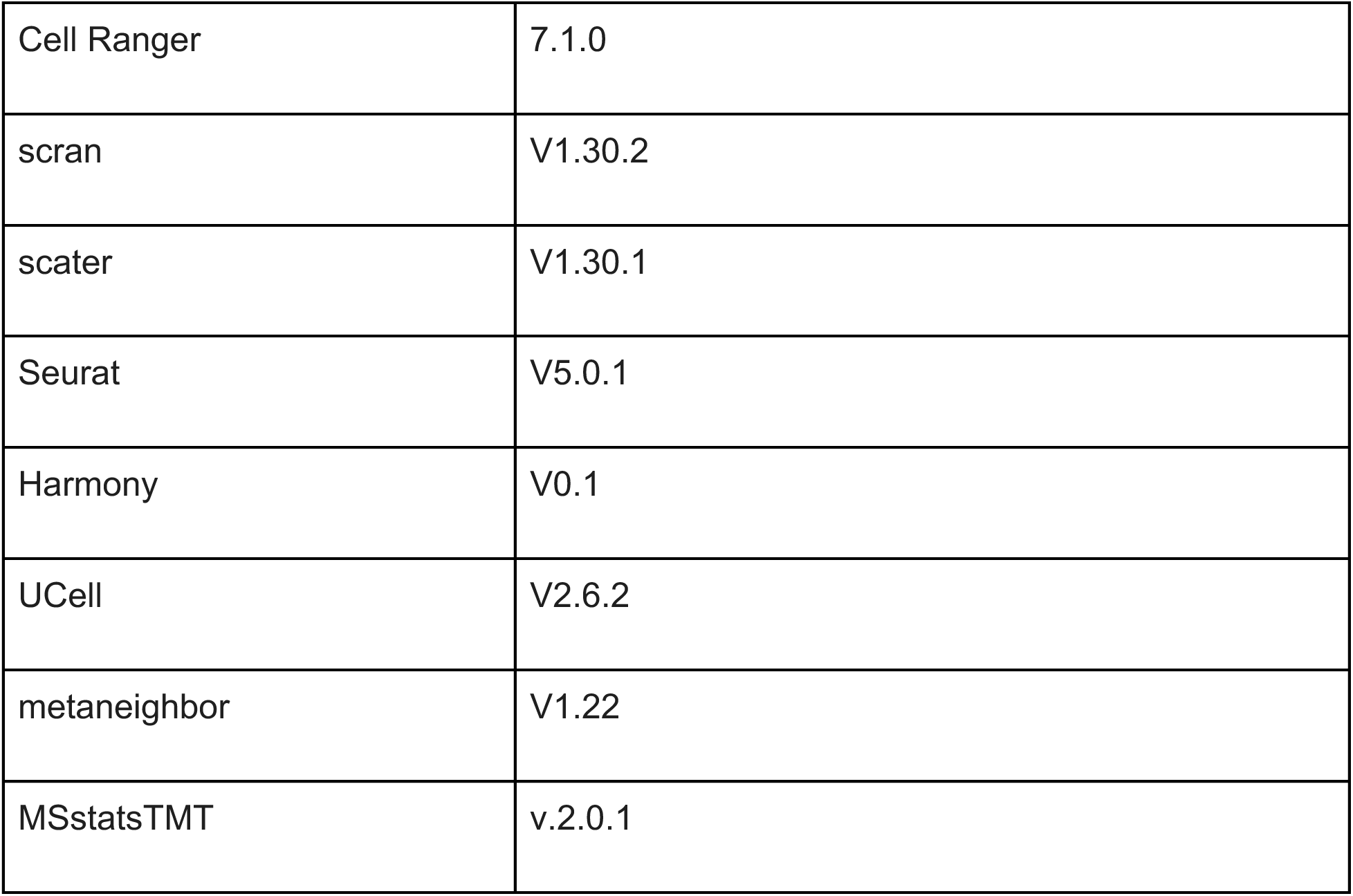
Software and Algorithms.

## Data availability

Sequencing data that support the findings of this study have been deposited in the Gene Expression Omnibus under the accession codes GSE298570 (RNA-seq data), GSE298571 (scRNA-seq data).

## Acknowledgments

The authors would like to acknowledge Baris Bingol, Irena Kadiu, Haiyang Yu and Maggie Crow for critical inputs of the manuscript; Brad Friedman for bioinformatics consultation; Zora Modrusan and the NGS core facility for performing RNA sequencing; Chris Bohlen for advice and consultation; Monica Xiong, Jessica Lawrence, Allison Soung and Sean Silverman for materials and knowledge sharing; Ana M Meireles and Sylvia Fechner for multi-omics insights and expertise; Alstem and Jackson Laboratory for iPS lines, Neuroscience department and Genentech for all the support.

