## Supplementary figures S1-S7 for "Integrated morphological, multi-omics, and functional profiling reveals microglial plasticity driven by AD risk genes"

Fig. S1

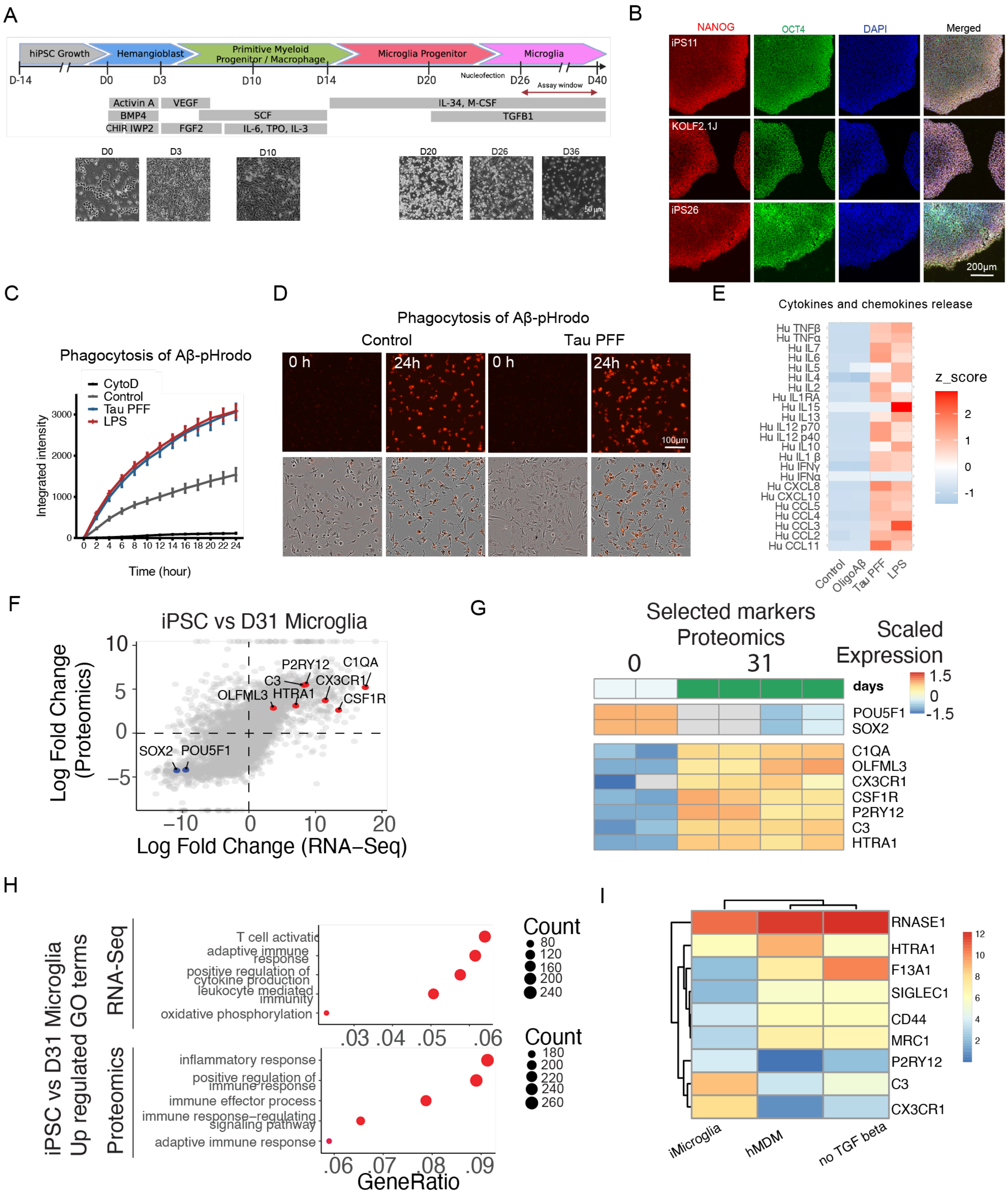
A
hiPSC Growth
Hemangioblast
Primitive Myeloid Progenitor / Macrophage
Microglia Progenitor
Microglia
D-14
D0
D3
D10
D14
D20
Nucleofection
D26
Assay window
D40
Activin A
BMP4
CHIR IWP2
VEGF
FGF2
SCF
IL-6, TPO, IL-3
IL-34, M-CSF
TGFB1
D0
D3
D10
D20
D26
D36
B
NANOG
OCT4
DAPI
Merged
iPS11
KOLF2.1J
iPS26
200μm
C
Phagocytosis of Aβ-pHrodo
CytoD
Control
Tau PFF
LPS
Integrated intensity
Time (hour)
D
Phagocytosis of Aβ-pHrodo
Control
Tau PFF
0 h
24h
100μm
E
Cytokines and chemokines release
Hu TNFβ
Hu TNFα
Hu IL7
Hu IL6
Hu IL5
Hu IL4
Hu IL2
Hu IL1RA
Hu IL15
Hu IL13
Hu IL12 p70
Hu IL12 p40
Hu IL10
Hu IL1 β
Hu IFNγ
Hu IFNα
Hu CXCL8
Hu CXCL10
Hu CCL5
Hu CCL4
Hu CCL3
Hu CCL2
Hu CCL11
z_score
Control
OligoAβ
Tau PFF
LPS
F
iPSC vs D31 Microglia
Log Fold Change (Proteomics)
Log Fold Change (RNA−Seq)
P2RY12
C1QA
C3
CX3CR1
OLFML3
HTRA1
CSF1R
SOX2
POU5F1
G
Selected markers Proteomics
0
31
Scaled Expression
days
POU5F1
SOX2
C1QA
OLFML3
CX3CR1
CSF1R
P2RY12
C3
HTRA1
H
iPSC vs D31 Microglia Up regulated GO terms
RNA-Seq
T cell activation
adaptive immune response
positive regulation of cytokine production
leukocyte mediated immunity
oxidative phosphorylation
Count
Proteomics
inflammatory response
positive regulation of immune response
immune effector process
immune response−regulating signaling pathway
adaptive immune response
Count
GeneRatio
I
RNASE1
HTRA1
F13A1
SIGLEC1
CD44
MRC1
P2RY12
C3
CX3CR1
iMicroglia
hMDM
no TGF beta

**Supplementary Figure 1. iPSC-derived microglia functionally and transcriptionally recapitulate key features of human microglia.**

**A.**Schematic illustrating the differentiation of iMicroglia from human iPSCs via primitive hematopoiesis. Representative time-course bright-field images depict iMicroglia differentiation from iPSCs, showing the distinct morphology and patterns at each stage.

**B.** Representative IF images showing the expression of pluripotent markers NANOG and OCT4 in the three iPS lines used in the study.

**C.** Time-course phagocytosis of Aβ-pHrodo by iMicroglia derived from iPS11. Engulfment of Aβ-pHrodo was completely inhibited with Cytocalasin D (CytoD) treatment. iMicroglia pretreated with Tau PFFs (10 µg/ml) or LPS (100 ng/ml) showed increased engulfment of Aβ-pHrodo.

**D.** Representative images showing phagocytosis of Aβ-pHrodo at 0 and 24 hours in untreated control and Tau PFF-treated iMicroglia. Increased accumulation of Aβ-pHrodo was observed in Tau PFF-treated groups.

**E.** Heatmap of average amount of cytokines and chemokines secreted by iMicroglia stimulated with oligomeric Aβ (1 µM), Tau PFFs (10 µg/mL) or LPS (25 ng/mL). The release of cytokines and chemokines were increased in Tau PFF or LPS - treated iMicroglia.

**F.** Correlation of RNA and protein log fold change between iPSC and iMicroglia was calculated (r = 0.664).

**G.** Heatmap generated from proteomics data showing mature microglia markers are expressed at the protein level in iMicroglia.

**H.** Gene ontology (GO) enrichment analysis showing significant enrichment of immune related terms at both RNA and protein levels (top 5 terms FDR < 0.05)

**I.** Heatmap analysis of RNA-seq data showed that iMicroglia highly expressed core microglia signature genes (P2RY12, C3, CX3CR1), while concurrently downregulating traditional macrophage markers (HTRA1, F13A1, SIGLEC1, etc). This expression pattern contrasts sharply with that observed in hBMDM and microglia cultures lacking TGF-β.

Fig. S2

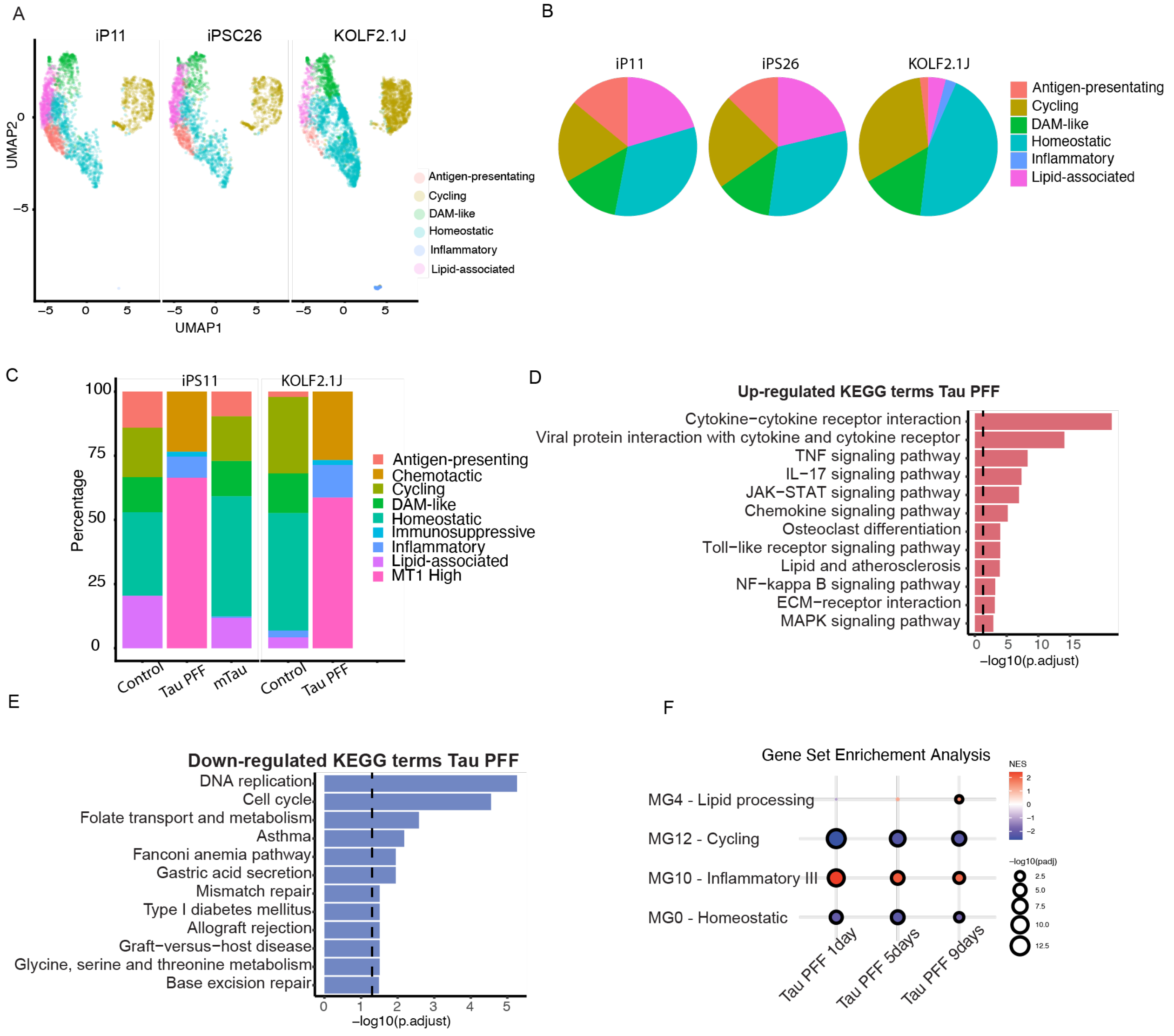
A
iP11
iPSC26
KOLF2.1J
UMAP2
UMAP1
Antigen-presentating
Cycling
DAM-like
Homeostatic
Inflammatory
Lipid-associated
B
iP11
iPS26
KOLF2.1J
Antigen-presentating
Cycling
DAM-like
Homeostatic
Inflammatory
Lipid-associated
C
iPS11
KOLF2.1J
Percentage
Control
Tau PFF
mTau
Control
Tau PFF
Antigen-presenting
Chemotactic
Cycling
DAM-like
Homeostatic
Immunosuppressive
Inflammatory
Lipid-associated
MT1 High
D
Up-regulated KEGG terms Tau PFF
Cytokine−cytokine receptor interaction
Viral protein interaction with cytokine and cytokine receptor
TNF signaling pathway
IL−17 signaling pathway
JAK−STAT signaling pathway
Chemokine signaling pathway
Osteoclast differentiation
Toll−like receptor signaling pathway
Lipid and atherosclerosis
NF−kappa B signaling pathway
ECM−receptor interaction
MAPK signaling pathway
−log10(p.adjust)
E
Down-regulated KEGG terms Tau PFF
DNA replication
Cell cycle
Folate transport and metabolism
Asthma
Fanconi anemia pathway
Gastric acid secretion
Mismatch repair
Type I diabetes mellitus
Allograft rejection
Graft−versus−host disease
Glycine, serine and threonine metabolism
Base excision repair
−log10(p.adjust)
F
Gene Set Enrichement Analysis
MG4 - Lipid processing
MG12 - Cycling
MG10 - Inflammatory III
MG0 - Homeostatic
Tau PFF 1day
Tau PFF 5days
Tau PFF 9days
NES
−log10(padj)

**Supplement Figure 2. Treatment with Tau PFFs drives iMicroglia into more inflammatory and oxidative stress–adaptive states marked by activation of TNF and NF-κB signaling.**

**A.** Single-cell UMAP of iMicroglia derived from iPS11, iPS26, and KOLF2.1J lines reveal comparable state compositions, demonstrating robust protocol consistency across iPSC lines.

**B**. Pie charts showing the distribution of microglial states across the three iPS-derived iMicroglia lines.

**C**.The proportions of different clusters are similar between control and Tau monomer-treated cells, whereas Tau PFF treatment shifts cells toward MT1-high, chemotactic, and inflammatory states.

**D-E**. KEGG pathway enrichment analysis of Tau PFF-treated cells reveals prominent upregulation of inflammatory-related pathways (D) and downregulation of cell-cycle pathways (E). Dotted line denotes FDR < 0.05.

**F**. GSEA of bulk RNA-seq data shows that genes upregulated at 9 days after Tau PFF treatment are significantly enriched for the lipid processing, DAM-like signature corresponding to the in vivo MG4 cluster defined by Sun et al. (dotted outlines indicate adjusted p value < 0.05).

Fig. S3

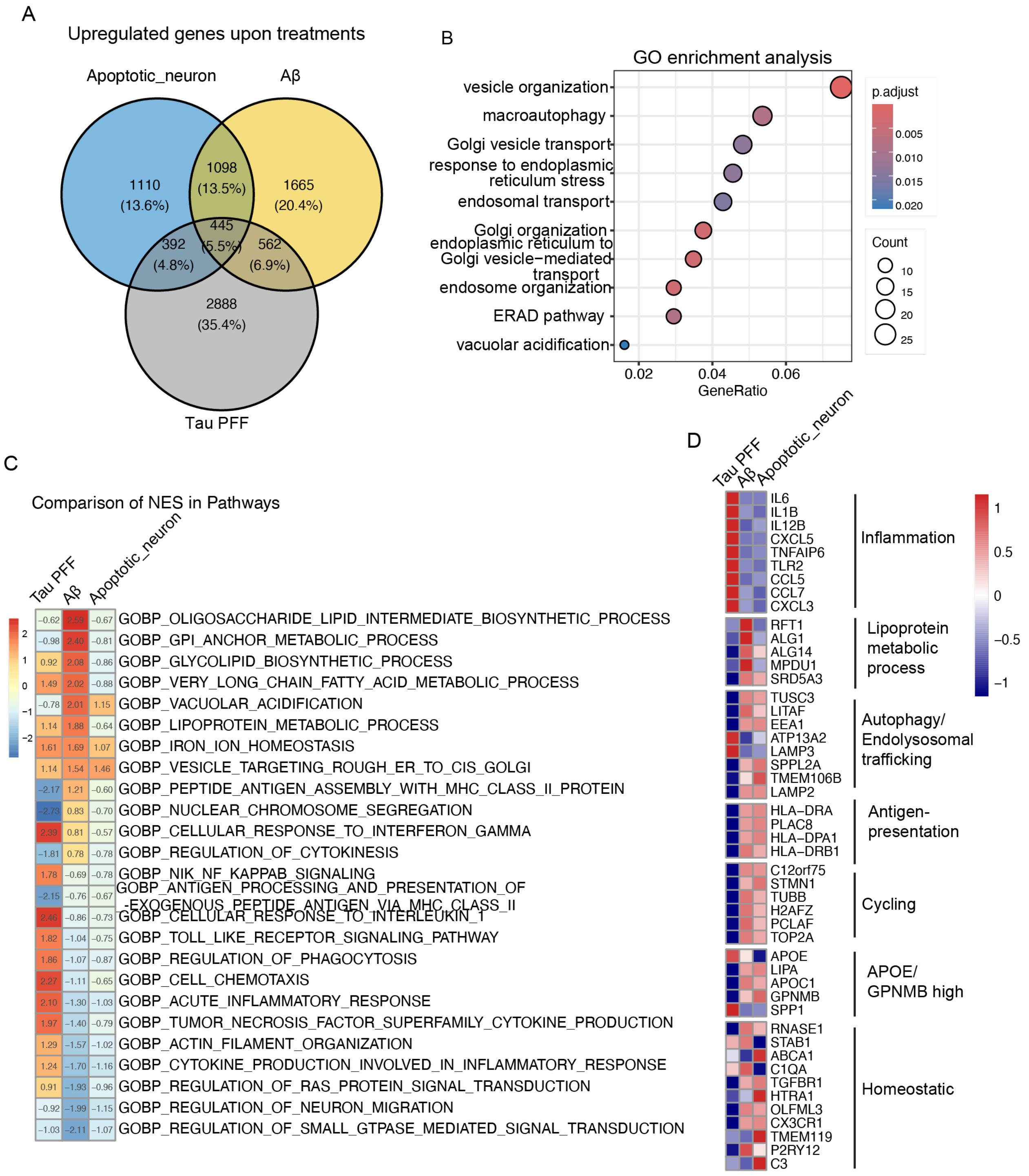
A
Upregulated genes upon treatments
Apoptotic_neuron
Aβ
Tau PFF
1110 (13.6%)
1098 (13.5%)
1665 (20.4%)
445 (5.5%)
392 (4.8%)
562 (6.9%)
2888 (35.4%)
B
GO enrichment analysis
vesicle organization
macroautophagy
Golgi vesicle transport
response to endoplasmic reticulum stress
endosomal transport
Golgi organization
endoplasmic reticulum to Golgi vesicle−mediated transport
endosome organization
ERAD pathway
vacuolar acidification
0.02 0.04 0.06
GeneRatio
p.adjust
0.005
0.010
0.015
0.020
Count
10
15
20
25
C
Comparison of NES in Pathways
Tau PFF
Aβ
Apoptotic_neuron
2
1
0
−1
−2
−0.62 2.59 −0.67 GOBP_OLIGOSACCHARIDE_LIPID_INTERMEDIATE_BIOSYNTHETIC_PROCESS
−0.98 2.40 −0.81 GOBP_GPI_ANCHOR_METABOLIC_PROCESS
0.92 2.08 −0.86 GOBP_GLYCOLIPID_BIOSYNTHETIC_PROCESS
1.49 2.02 −0.88 GOBP_VERY_LONG_CHAIN_FATTY_ACID_METABOLIC_PROCESS
−0.78 2.01 1.15 GOBP_VACUOLAR_ACIDIFICATION
1.14 1.88 −0.64 GOBP_LIPOPROTEIN_METABOLIC_PROCESS
1.61 1.69 1.07 GOBP_IRON_ION_HOMEOSTASIS
1.14 1.54 1.46 GOBP_VESICLE_TARGETING_ROUGH_ER_TO_CIS_GOLGI
−2.17 1.21 −0.60 GOBP_PEPTIDE_ANTIGEN_ASSEMBLY_WITH_MHC_CLASS_II_PROTEIN
−2.73 0.83 −0.70 GOBP_NUCLEAR_CHROMOSOME_SEGREGATION
2.39 0.81 −0.57 GOBP_CELLULAR_RESPONSE_TO_INTERFERON_GAMMA
−1.81 0.78 −0.78 GOBP_REGULATION_OF_CYTOKINESIS
1.78 −0.69 −0.78 GOBP_NIK_NF_KAPPAB_SIGNALING
−2.15 −0.76 −0.67 GOBP_ANTIGEN_PROCESSING_AND_PRESENTATION_OF_EXOGENOUS_PEPTIDE_ANTIGEN_VIA_MHC_CLASS_II
2.46 −0.86 −0.73 GOBP_CELLULAR_RESPONSE_TO_INTERLEUKIN_1
1.82 −1.04 −0.75 GOBP_TOLL_LIKE_RECEPTOR_SIGNALING_PATHWAY
1.86 −1.07 −0.87 GOBP_REGULATION_OF_PHAGOCYTOSIS
2.27 −1.11 −0.65 GOBP_CELL_CHEMOTAXIS
2.10 −1.30 −1.03 GOBP_ACUTE_INFLAMMATORY_RESPONSE
1.97 −1.40 −0.79 GOBP_TUMOR_NECROSIS_FACTOR_SUPERFAMILY_CYTOKINE_PRODUCTION
1.29 −1.57 −1.02 GOBP_ACTIN_FILAMENT_ORGANIZATION
1.24 −1.70 −1.16 GOBP_CYTOKINE_PRODUCTION_INVOLVED_IN_INFLAMMATORY_RESPONSE
0.91 −1.93 −0.96 GOBP_REGULATION_OF_RAS_PROTEIN_SIGNAL_TRANSDUCTION
−0.92 −1.99 −1.15 GOBP_REGULATION_OF_NEURON_MIGRATION
−1.03 −2.11 −1.07 GOBP_REGULATION_OF_SMALL_GTPASE_MEDIATED_SIGNAL_TRANSDUCTION
D
Tau PFF
Aβ
Apoptotic_neuron
IL6
IL1B
IL12B
CXCL5
TNFAIP6
TLR2
CCL5
CCL7
CXCL3
Inflammation
RFT1
ALG1
ALG14
MPDU1
SRD5A3
Lipoprotein metabolic process
TUSC3
LITAF
EEA1
ATP13A2
LAMP3
SPPL2A
TMEM106B
LAMP2
Autophagy/ Endolysosomal trafficking
HLA−DRA
PLAC8
HLA−DPA1
HLA−DRB1
Antigen-presentation
C12orf75
STMN1
TUBB
H2AFZ
PCLAF
TOP2A
Cycling
APOE
LIPA
APOC1
GPNMB
SPP1
APOE/ GPNMB high
RNASE1
STAB1
ABCA1
C1QA
TGFBR1
HTRA1
OLFML3
CX3CR1
TMEM119
P2RY12
C3
Homeostatic
1
0.5
0
−0.5
−1

**Supplementary Figure 3. Specific disease-relevant stimuli, Tau and Aβ PFFs, induce distinct transcriptional responses in iMicroglia, suggesting that each triggers unique signaling pathways.**

**A**. Venn diagram illustrating the shared upregulated genes among iMicroglia exposed to Tau PFFs, Aβ fibrils, and apoptotic neurons. The intersection reveals 445 upregulated genes in all three stimuli.

**B**. GO analysis of the 445 shared upregulated genes (Supplementary Fig. 4A) illustrated significant enrichment in vesicle organization and trafficking pathways.

**C**. Heatmap comparing the normalized enrichment scores (NES) of select significantly enriched pathways identified from GSEA of cells treated with Tau PFFs, Aβ, or apoptotic neurons.

**D**. Heatmap illustrates gene expression patterns associated with relevant biological processes and cell states. Tau PFF treatment induced the upregulation of inflammatory-related genes and the downregulation of genes marking antigen presentation, cell cycling, and homeostatic states. Aβ treatment significantly increased the expression of genes involved in lipoprotein metabolic processes.

Fig. S4

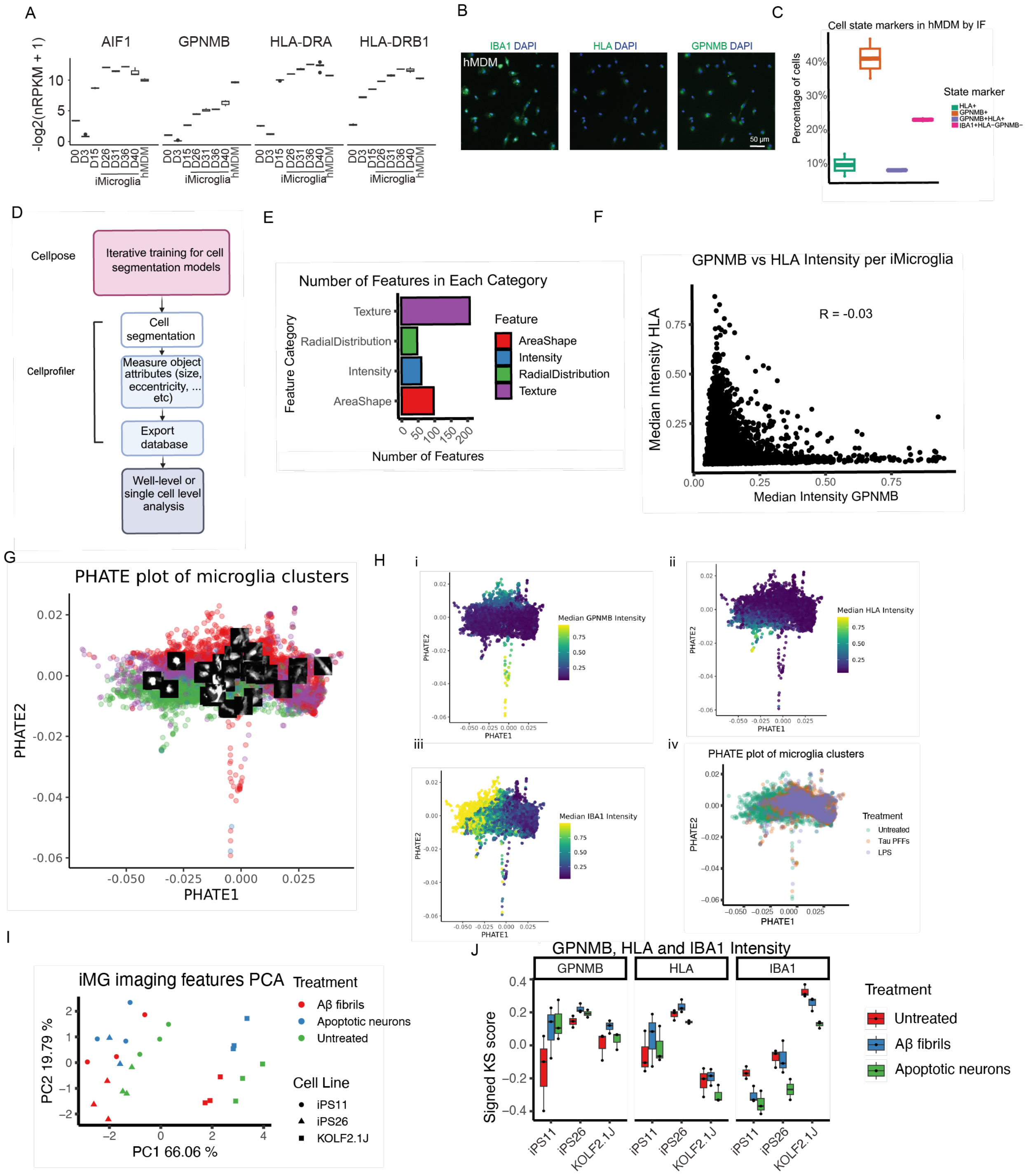

A
AIF1
GPNMB
HLA-DRA
HLA-DRB1
-log2(nRPKM + 1)
iMicroglia
hMDM
B
IBA1 DAPI
HLA DAPI
GPNMB DAPI
hMDM
50 µm
C
Cell state markers in hMDM by IF
Percentage of cells
State marker
HLA+
GPNMB+
GPNMB+HLA+
IBA1+HLA-GPNMB-
D
Cellpose
Iterative training for cell segmentation models
Cellprofiler
Cell segmentation
Measure object attributes (size, eccentricity, ... etc)
Export database
Well-level or single cell level analysis
E
Number of Features in Each Category
Feature Category
Texture
RadialDistribution
Intensity
AreaShape
Number of Features
Feature
AreaShape
Intensity
RadialDistribution
Texture
F
GPNMB vs HLA Intensity per iMicroglia
R = -0.03
Median Intensity HLA
Median Intensity GPNMB
G
PHATE plot of microglia clusters
PHATE2
PHATE1
H
Median GPNMB Intensity
Median HLA Intensity
Median IBA1 Intensity
PHATE plot of microglia clusters
Treatment
Untreated
Tau PFFs
LPS
I
iMG imaging features PCA
Treatment
Aβ fibrils
Apoptotic neurons
Untreated
Cell Line
iPS11
iPS26
KOLF2.1J
PC1 66.06 %
PC2 19.79 %
J
GPNMB, HLA and IBA1 Intensity
GPNMB
HLA
IBA1
Signed KS score
Treatment
Untreated
Aβ fibrils
Apoptotic neurons

**Supplemental Figure 4. Microglial transcriptional states are substantiated by IF and map to distinct morphological signatures.**

**A.** RNA-seq analysis shows that expression of microglia markers (*AIF1*/*IBA1*) and antigen-presenting state markers (*HLA-DRA*, *HLA-DRB1*) is lower in hMDM compared to iMicroglia, whereas *GPNMB*, a DAM-like cell marker, is expressed at higher levels in hMDM cultures than in iMicroglia.

**B.** iMicroglia state markers were also applied to hMDM. Representative IF images of anti-IBA1, HLA and GPNMB signals are shown on hMDM.

**C.** Barplot showed the composition of clusters in hMDM (HLA+, GPNMB+, HLA+GPNMB+, IBA1+HLA-GPNMB- populations). GPNMB+ population is the predominant cluster in hMDM culture.

**D**. Image analysis framework. Segmentation models were trained using Cellpose2 to recognize IBA1+, GPNMB+ or HLA+ cells. These models were subsequently applied in Cellprofiler to segment the corresponding objects. Object relationship and quantitative features related to size and shape, intensity, intensity distribution and texture were extracted using Cellprofiler and analyzed at both the well level and the single cell level.

**E.** Barplot showing that the features extracted via CellProfiler can be categorized into texture, area shape, intensity and radial distribution. Numbers of features in each category are shown here.

**F.** Plot showed GPNMB intensity and HLA intensity of each iMicroglia cell is not correlated (R = -0.03). High GPNMB intensity cells tend to have low HLA intensity, and vice versa.

**G.** Dimensionality reduction was performed using the PHATE algorithm on iMicroglia's morphological features. Colors indicate cell states as determined by the Cellpose models. Ten IBA1+ objects were randomly selected from each state and plotted according to their perspective location in the Phate morphological feature space.

**H**. Median GPNMB(i), HLA(ii), IBA1 intensities (iii) intensities for each cell and treatment map (vi) are overlaid within the same Phate space in (G). Untreated and Tau PFF or LPS treated cells were separated along PHATE1. IBA1, HLA and GPNMB intensities were decreased with Tau PFF and LPS treatment.

**I.** PCA of iMicroglia morphological features treated with fibrillar Aβ and apoptotic neurons (AN) at the well level. Shapes indicate cell lines and colors indicate treatments. Both treatments shifted the morphology on PC1.

**J.** GPNMB median intensity is slightly increased by Aβ and AN treatment, while IBA1 median intensity is slightly decreased with both treatments in all three lines.

Fig. S5

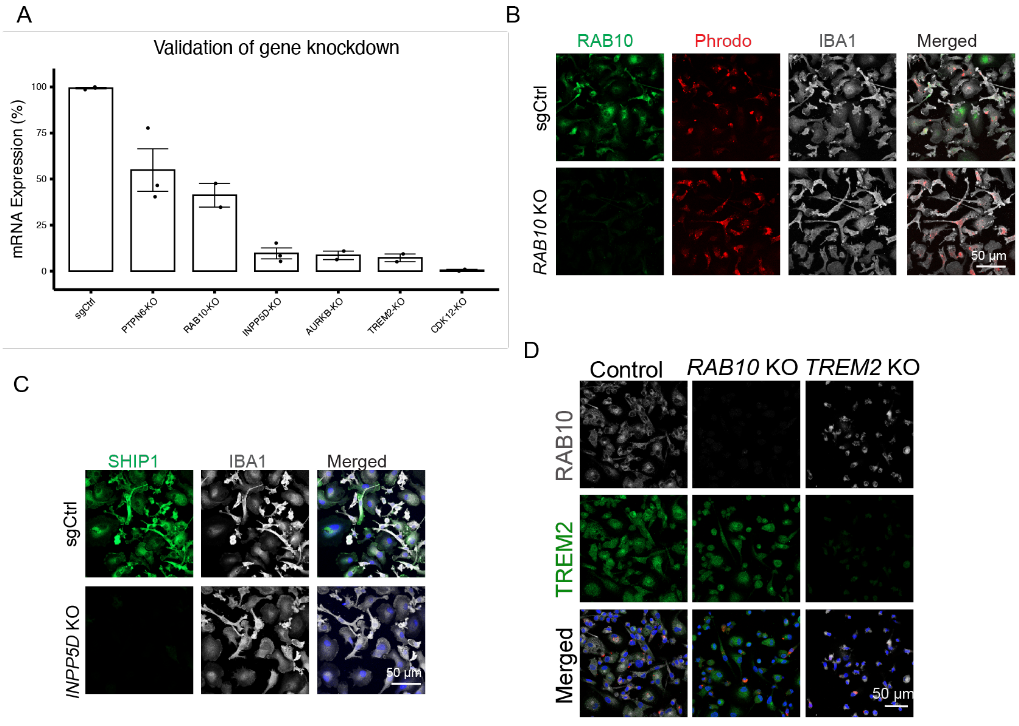


**Supplementary Figure 5. Validation of CRISPR knockout of AD GWAS genes in iMicroglia**

**A.** Bar plots showing mRNA levels of the targeted genes in iMicroglia following Cas9 RNP delivery, as measured by RT-PCR. Data are normalized to GAPDH and sgCtrl and presented as mean +/- SEM (n=3). Black dots indicate data distribution.

**B-D**. IF imaging showed protein RAB10, SHIP1 (encoded by *INPP5D*) and TREM2 were completely depleted upon CRISPR knockout.

Fig. S6

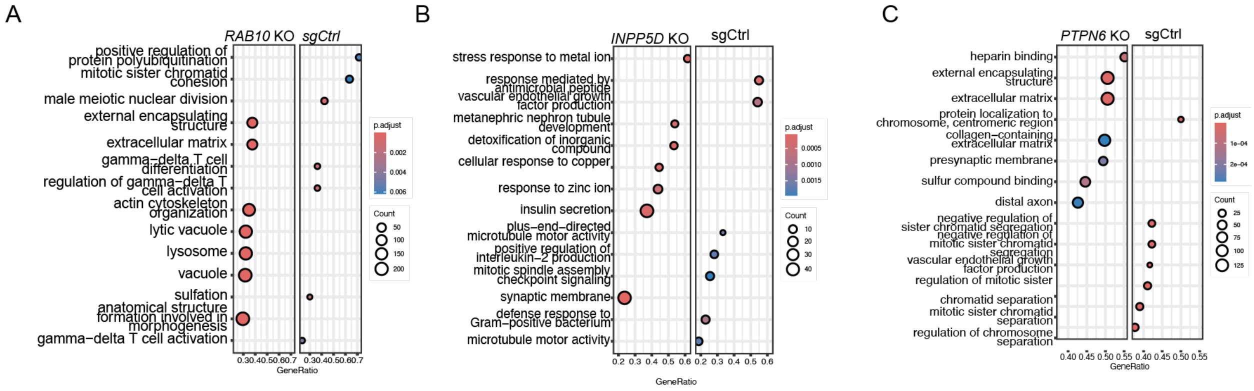


**Supplementary Figure 6. CRISPR-Cas9 KO of AD GWAS genes drives transcriptional changes**

**A.** Dot plot of top GSEA results for GO terms from differential expression analysis of control versus *RAB10* knockout iMicroglia. Dot size represents gene count, and dot color indicates Benjamini-Hochberg adjusted p-values. Lysosomal pathways are enriched in *RAB10* knockout cells.

**B-C.** Dot plot showing top results from GSEA of GO terms from differential expression analysis of control sgRNA vs *INPP5D* KO (B) and *PTPN6* KO (C) iMicroglia. Size of the dots represent the number of genes contributing to the GO terms. Color of dots represent adjusted statistical significance to account for multiple hypothesis testing using the Benjamini-Hochberg method. Upregulated genes in *INPP5D* KO iMicroglia are enriched for response to metal ion and detoxification processes and upregulated genes in *PTPN6* KO iMicroglia are enriched for extracellular matrix related processes.

Fig. S7

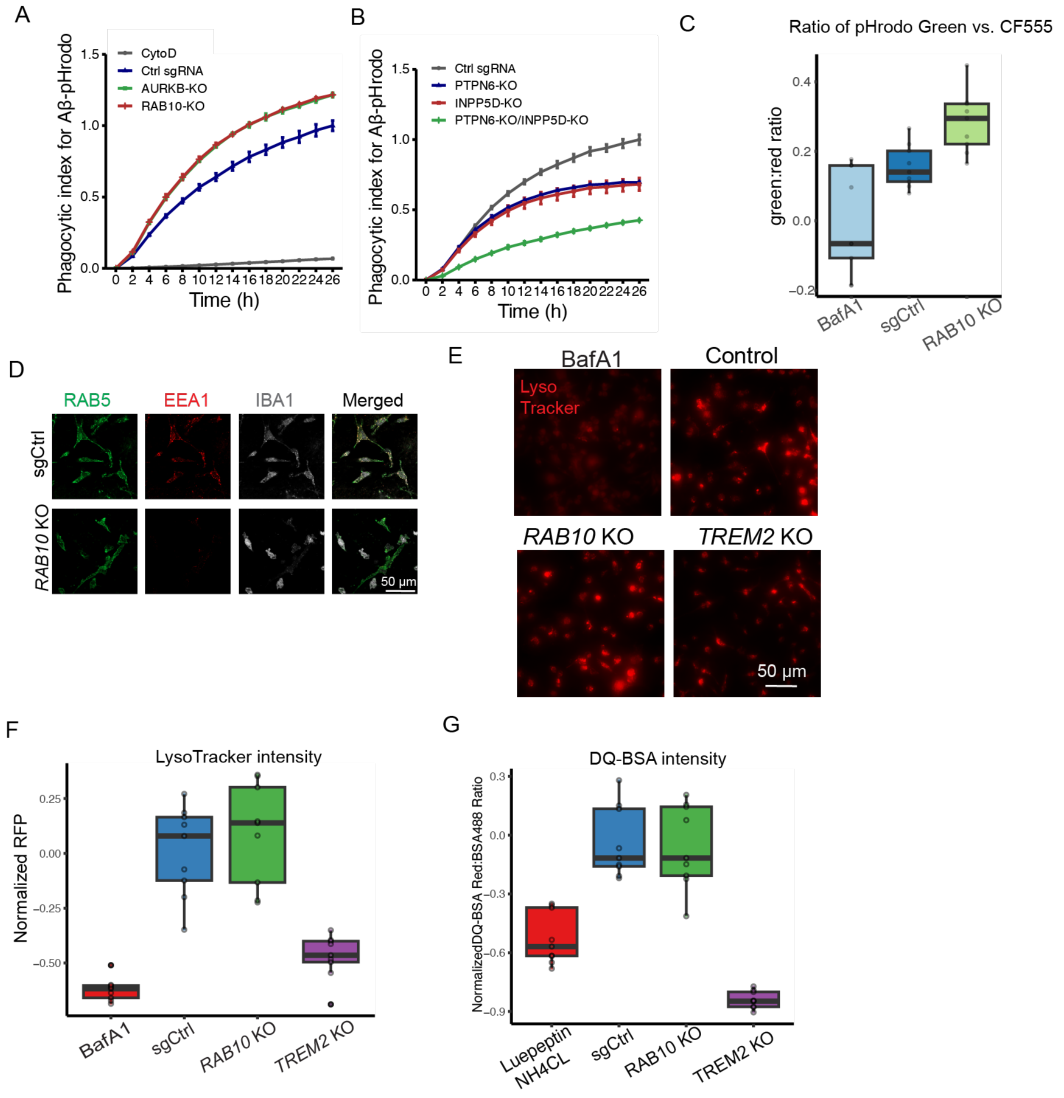


**Supplementary Figure 7. *RAB10* knockout disrupts endo-lysosomal homeostasis in iMicroglia.**

**A-B.** Time-course analysis of the phagocytic index for Aβ–pHrodo uptake in iMicroglia shows that depletion of *AURKB* or *RAB10* increases intracellular Aβ–pHrodo intensity. In contrast, depletion of *INPP5D* or *PTPN6* reduces Aβ–pHrodo intensity, with co-depletion resulting in a further decrease in cellular Aβ–pHrodo levels. iMicroglia treated with CytoD were used as a negative control. Data are presented as mean ± SEM (n = 3).

**C.** Boxplot showed the quantification of the ratio of pHrodo-green and CF555 intensity from three replicate experiments in Fig. 6E (n = 9 regions). The horizontal black lines within the boxes denote median values (50th percentiles), the black boxes contain the values from the 25th to the 75th percentiles and the black whiskers denote values at the 5th and 95th percentiles.The black circles represent the data distribution.

**D.** IF imaging showed that the expression of early endosomal marker EEA1 was reduced in *RAB10* depleted iMicroglia. Scale bar, 50 μm.

**E.** Representative images of control, *RAB10* KO and *TREM2* KO iMicroglia treated with LysoTracker Red dye (n = 9 images) (F). Cells were incubated with 1 μM BafA1 for 3 hours as a positive control. Depleting *RAB10* and *TREM2* both caused lysosomal deacidification.

**F.** Quantification of the normalized LysoTracker intensity per image (n= 9 images) in E.

**G**. Lysosomal hydrolytic activity determined by DQ$^{TM}$-BSA red/BSA-488 in iMicroglia expression control, *RAB10* and *TREM2* sgRNAs (n = 9 images). Leupeptin and $NH_4CL$-treated control iMicroglia served as a positive control. *TREM2* depletion caused impairment in hydrolytic activity.
